# Genome sequencing of ‘Fuji’ apple clonal varieties reveals evolutionary history and genetic mechanism of the spur-type morphology

**DOI:** 10.1101/2024.05.16.594617

**Authors:** Yudong Cai, Xiuhua Gao, Jiangping Mao, Yu Liu, Lu Tong, Xilong Chen, Yandong Liu, Wenyan Kou, Chuanjun Chang, Toshi Foster, Jialong Yao, Amandine Cornille, Muhammad Mobeen Tahir, Zhi Liu, Zhongye Yan, Siyi Lin, Fengwang Ma, Juanjuan Ma, Libo Xing, Na An, Xiya Zuo, Yanrong Lv, Zhengyang Zhao, Wenqiang Li, Qianjin Li, Caiping Zhao, Yanan Hu, Hangkong Liu, Chao Wang, Xueyan Shi, Doudou Ma, Zhangjun Fei, Yu Jiang, Dong Zhang

## Abstract

Somatic variations arising during asexual reproduction can lead to the emergence of bud sports exhibiting advantageous traits, forming the basis for bud sport breeding in perennial plants. Here, we report a near-complete, fully phased genome assembly of ‘Fuji’ apple, enabling comprehensive identification of somatic variants across 74 clonally propagated ‘Fuji’ varieties. Phylogenetic analysis of ‘Fuji’ sport varieties indicates that the emergence of spur-type and early maturation traits results from multiple independent formation events. A number of putative functional somatic variants are identified, including one spur-type specific deletion located in the promoter of the TCP transcription factor gene *MdTCP11*. DNA methylation level of the deletion-associated miniature inverted-repeat transposable element is lower in the spur-type varieties compared to the standard-type varieties, while the expression of *MdTCP11* is significantly higher. Overexpression of *MdTCP11* in apple decreases the plant height, highlighting its important role in the development of spur-type apple varieties. This study sheds light on the 80-year cloning history of ‘Fuji’ and provides valuable candidate causative mutations for apple breeding.

## Introduction

Somatic mutations occur in all living organisms, but studies on these mutations have largely focused on mosaic organs in the contexts of human aging, cancer, and neurodegeneration. In plants, somatic variations refer to the genetic changes that arise during the plant mitotic cell cycle. They are widespread in perennials such as fruit tree crops and contribute to genetic diversity, giving rise to new traits known as bud sports. Bud sports are the main source of new cultivars widely employed in apples, citruses, pears, and other fruit trees^1^. Vegetative propagation of bud sports enables the selection and breeding of clonal varieties; therefore, somatic variations can be preserved over long periods of time. The ability to clonally propagate mosaic or homogeneous somatic mutations is not only useful as a new source of genetic diversity but also allows us to identify the underlying mutations responsible for novel traits. However, high heterozygosity of the apple genome poses challenges in accurately identifying somatic mutations, as many heterozygous germline variants could be classified as somatic variants. Furthermore, structural variations or rearrangements between the two haploid genomes may complicate the detection of somatic mutations.

In China, 33.9% of apple cultivars are selected and bred through bud sport breeding^2^. ‘Fuji’ is one of the most prominent cultivars used in apple bud sport breeding, and numerous clonally propagated varieties have been selected from ‘Fuji’ through successive generations of bud sport breeding. Approximately 73.6% of cultivars planted in China are ‘Fuji’ bud sport clones with a wide range of new traits, such as spur-type growth habit, red peel and early maturation. The primary apple-producing regions in China are characterized by poor soil quality and frequent drought conditions, coupled with a scarcity of suitable dwarf rootstocks. The widespread adoption of spur-type varieties in high-density cultivation practices, driven by the agricultural conditions in China, significantly enhances apple productivity and orchard establishment. Spur-type apple trees have more concentrated flower bud differentiation and spur formation, which improves fruit yield and reduces pruning costs. Spur-type varieties are characterized by their more compact tree architecture and reduced tree size compared to the more open canopy form of standard-type varieties^3^. With shorter and thicker internodes, they produce shorter branches that are more suitable for pruning and fruit harvesting^4,5^. Previous studies have revealed that somatic mutations are typically observed as bud sports in apples, pears, citruses, and grapes^6–10^, and whole genome sequencing has been used to identify somatic variants^6,11^. In addition, some structural variants (SVs) associated with transposable element (TE) activities are also reported as a factor in bud sports^12–14^. Although multiple studies have analyzed the formation mechanism of spur-type varieties from the perspectives of endogenous hormones, transcriptomes and epigenetics^4,15,16^, the genomic basis of spur-type bud sports still remains unclear.

In this study, we have assembled a near-complete, fully phased diploid genome of ‘Fuji’. Using this diploid genome as a reference, we detect high-confidence somatic variants from 74 clonally propagated varieties of ‘Fuji’ and illuminate major candidate variants and underlying regulatory mechanisms for the spur-type morphology of apples. The study lays a foundation for studying the mechanisms of bud-sport emergence and the causative variants of bud sports, and provides valuable resources and knowledge for apple breeding.

## Results

### Phased diploid genome assembly of ‘Fuji’

‘Fuji’ was bred in 1939 through a cross between ‘Delicious’ and ‘Ralls Janet’. To assemble a high-quality ‘Fuji’ genome, we generated 101.36 Gb of PacBio HiFi sequencing data for four and 404.65 Gb of ultra-long Oxford Nanopore Technology (ONT) reads for 13 ‘Fuji’ clonally propagated varieties (**Supplementary Table 1**). To distinguish different haplotypes inherited from the two parents, we also generated 239.77 Gb of short-read data for the two parents, ‘Delicious’ and ‘Ralls Janet’ (**Supplementary Table 1**). Using the trio-binning approach, we first generated four haplotype-resolved contig assemblies for four varieties (‘Nagafu No.2’, ‘Fengfu2021’, ‘Red general’, ‘Red general spur’) with HiFi reads. The ultra-long ONT reads longer than 50 kb (48.45 Gb in total) were integrated into this process to improve the continuity of contigs. Using the assembly with the best continuity (‘Nagafu No.2’), we first employed a reference-guided method for scaffolding, and then generated 78.43 Gb Hi-C reads for ‘Nagafu No.2’, which were used to guide the manual correction of potential misassemblies (**Supplementary Fig. 1**). Finally, we used haplotype-resolved contigs from all four varieties to patch the gaps in the two haploid assemblies, resulting in a near-complete, fully phased diploid genome of ‘Fuji’ (**Fig. 1a**).

**Fig. 1.**
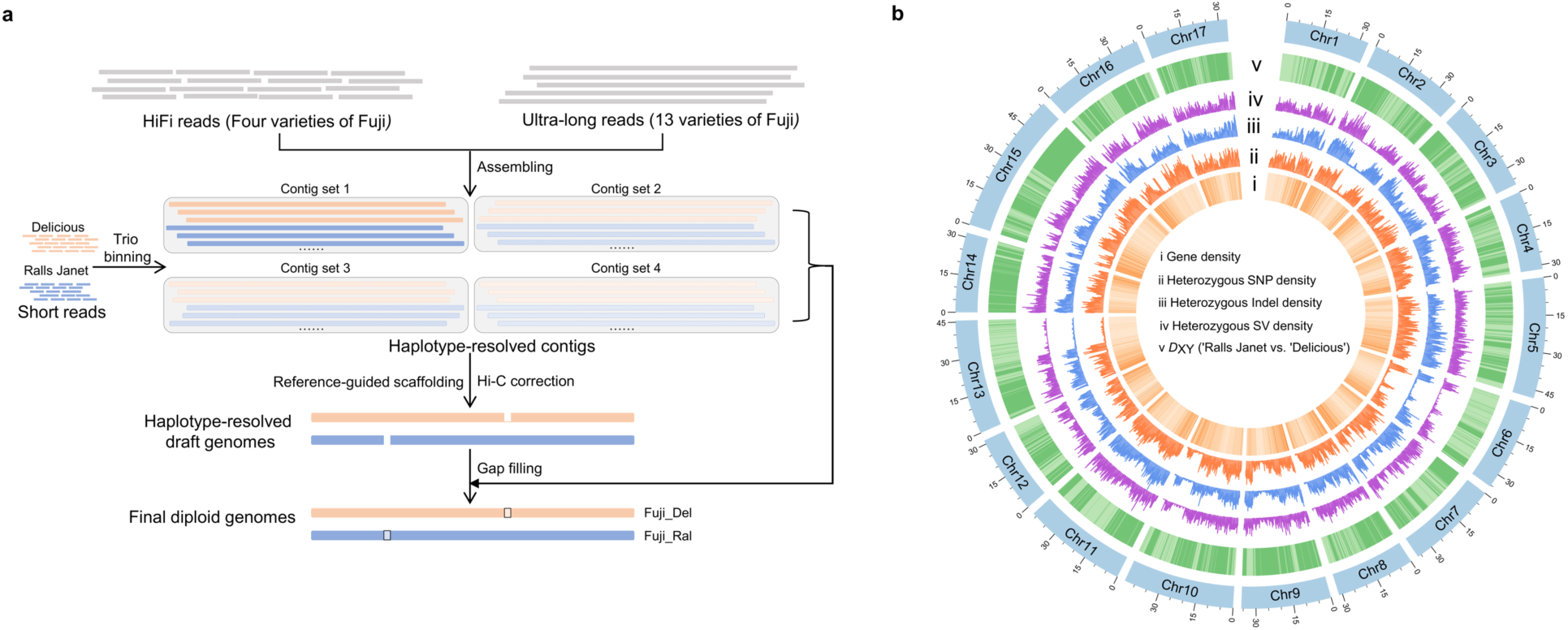
Genomic features of ‘Fuji’. a. Assembly strategy of the ‘Fuji’ diploid genome. The HiFi reads of four ‘Fuji’ clonally propagated varieties were assembled into contigs, integrated with the ultra-long reads of 13 other ‘Fuji’ clonally propagated varieties. Short reads from ‘Fuji’ parents, ‘Delicious’ and ‘Ralls Janet’, were used for phasing. **b** Genomic features of the ‘Fuji’ genome assembly (window size of 0.1Mb). *D*_XY_, absolute genetic divergence.

The phased assembly of ‘Fuji’ contained two haploid genomes (haplomes), one inherited from ‘Ralls Janet’ (hereafter Fuji_Ral) and the other from ‘Delicious’ (hereafter Fuji_Del). The assembly sizes of the two haplomes were 661.58 Mb and 658.72 Mb, respectively, similar to those of the previously published apple genome assemblies^14,16–18^. The total lengths of the 17 nuclear chromosomes of Fuji_Del and Fuji_Ral were 654.62 Mb and 653.37 Mb, respectively, and 14 chromosomes had paired telomeres in at least one haplome, and the remaining three chromosomes had only single telomeres (**Supplementary Table 2**). The two haplomes showed a high level of similarity (**Supplementary Fig. 2**) and had a high degree of collinearity with the GDDH13 genome^16^ (**Supplementary Fig. 3).**

We evaluated the quality of Fuji_Ral and Fuji_Del using multiple methods. The contig N50 values of the two haploid genomes were 37.30 Mb and 36.89 Mb, respectively, which were higher than those of the previously published apple genomes (**Supplementary Table 2**). The LTR assembly index (LAI) was 20.73 and 21.34 for Fuji_Ral and Fuji_Del, reaching the ‘gold standard’^19^. The BUSCO^20^ completeness rates of Fuji_Ral and Fuji_Del were 98.3% and 98.8%, respectively. Furthermore, the two haploid genomes reached a high quality of consensus base call, with the quality value (QV) of 59.06 for Fuji_Ral and 58.70 for Fuji_Del, close to the level of QV60 in recent high-quality diploid assemblies^21–23^.

We predicted a total of 50,646 and 50,678 genes in Fuji_Ral and Fuji_Del, respectively, and most of the homologous genes between ‘Fuji’ and GDDH13 had a high similarity (median 99.60%) (**Supplementary Fig. 4**). We annotated 458.75 Mb and 456.80 Mb of repetitive sequences that accounted for 69.34% and 69.35% of Fuji_Ral and Fuji_Del, respectively. The repeated sequence composition of the two haploid genomes was similar (**Supplementary Fig. 5**). We detected sequences of insertions and deletions between the two haploid genomes of ‘Fuji’, uncovering haplotype-specific sequences amounting to 73.87 Mb. This observation was further validated by genomic data across different ‘Fuji’ varieties, indicating that these sequences likely represent the hemizygous regions within the ‘Fuji’ genome (**Supplementary Table 3**).

To accurately identify genetic differences between the two haplomes in the ‘Fuji’ cultivar population, we further sequenced 74 clonally propagated varieties of ‘Fuji’, obtaining a total of 2,215.10 Gb of Illumina sequences (**Supplementary Table 4**). Through both genome alignment and read-based methods at the population level, we detected 4,478,495 single-nucleotide polymorphisms (SNPs), 480,098 small insertions/deletions (indels), and 16,734 SVs between the two ‘Fuji’ haplomes (**Supplementary Tables 5 and 6**). These variants represented heterozygous alleles shared among different ‘Fuji’ varieties inherited from the common ancestor (referred to as germline variants). Interestingly, we observed an unequal distribution of these heterozygous regions across the genome (**Fig. 1b**). We scanned the nucleotide divergence between ‘Ralls Janet’ and ‘Delicious’, the two parents of ‘Fuji’, along the ‘Fuji’ genome, and found that regions with low divergence coincided with those with low heterozygosity in the ‘Fuji’ genome (**Fig. 1b**), in line with the formation history of ‘Fuji’.

### Somatic variations in ‘Fuji’ clonally propagated varieties

During the clonal propagation of ‘Fuji’, different varieties have been selected based on their traits caused by somatic variations. However, in the context of the high heterozygosity of the ‘Fuji’ genome, there are much more heterozygous germline variants that differ between the two haploid genomes than somatic variants, which may lead to numerous germline variants being misidentified as somatic variants. Therefore, to eliminate germline variants as many as possible in detecting somatic variants, we used the phased diploid ‘Fuji’ genome as the reference by assigning each read to the appropriate haplomes (**Fig. 2a**). We detected a total of 68,965 somatic SNPs in the 74 individuals, including 532 deleterious mutations (SIFT score ≤ 0.05). Furthermore, we also identified 27,757 somatic indels and 1,848 somatic SVs (**Supplementary Tables 7-9**).

**Fig. 2.**
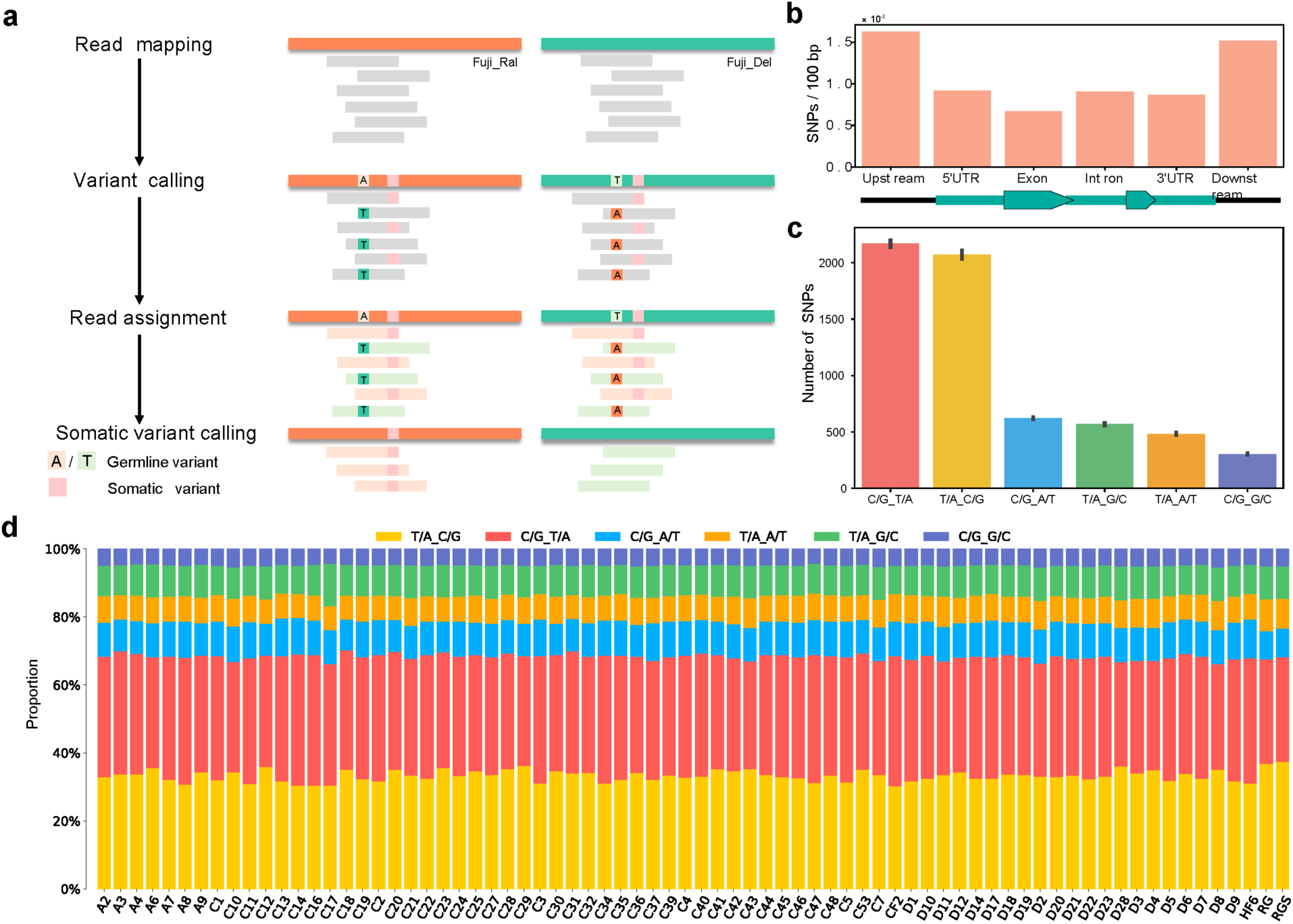
Somatic variants in ‘Fuji’ clonally propagated varieties. a. Pipeline of somatic variant detection. Each read is aligned to Fuji_Ral and Fuji_Del, respectively, and then assigned to the optimal haploid genome based on the alignment quality. **b** Densities of somatic SNPs in different gene features. **c** Number of different types of somatic SNPs in the population. **d** Distribution of different types of somatic SNPs in each variety.

Among the somatic SNPs, the proportion of the two transition types was higher than that of the four transversion types (**Fig. 2c**). Among them, the frequencies of C->T and C->A mutations were higher than those of the other transition and transversion types, respectively (**Fig. 2c**). This bias of mutant types is also widely observed in other plant species, consistent with the expected signatures of somatic variants^24–26^. We calculated the density of somatic SNPs in different genomic regions and found that somatic variants showed a similar pattern to germline variants (**Fig. 2b** and **Supplementary Fig. 6**), with the highest density found in the upstream and downstream regions of genes, followed by 5’ UTR, 3’ UTR and introns. In contrast, the lowest density was observed in exons, consistent with the conserved nature of exons.

### Population structure of ‘Fuji’ varieties

We performed principal component analysis (PCA) using all germline and somatic SNPs detected based on one haploid genome (Fuji_Ral) to evaluate the genetic relationship among different cultivars. The first and second principal components clearly distinguished the three groups: ‘Fuji’, ‘Ralls Janet’, and ‘Delicious’ (**Fig. 3a**). We further confirmed the genetic relationships among and within these three groups by kinship analysis, which revealed that the 74 ‘Fuji’ varieties were clonally related and were the offspring of ‘Ralls Janet’ and ‘Delicious’, consistent with the known breeding history of ‘Fuji’ (**Fig. 3b**).

**Fig. 3.**
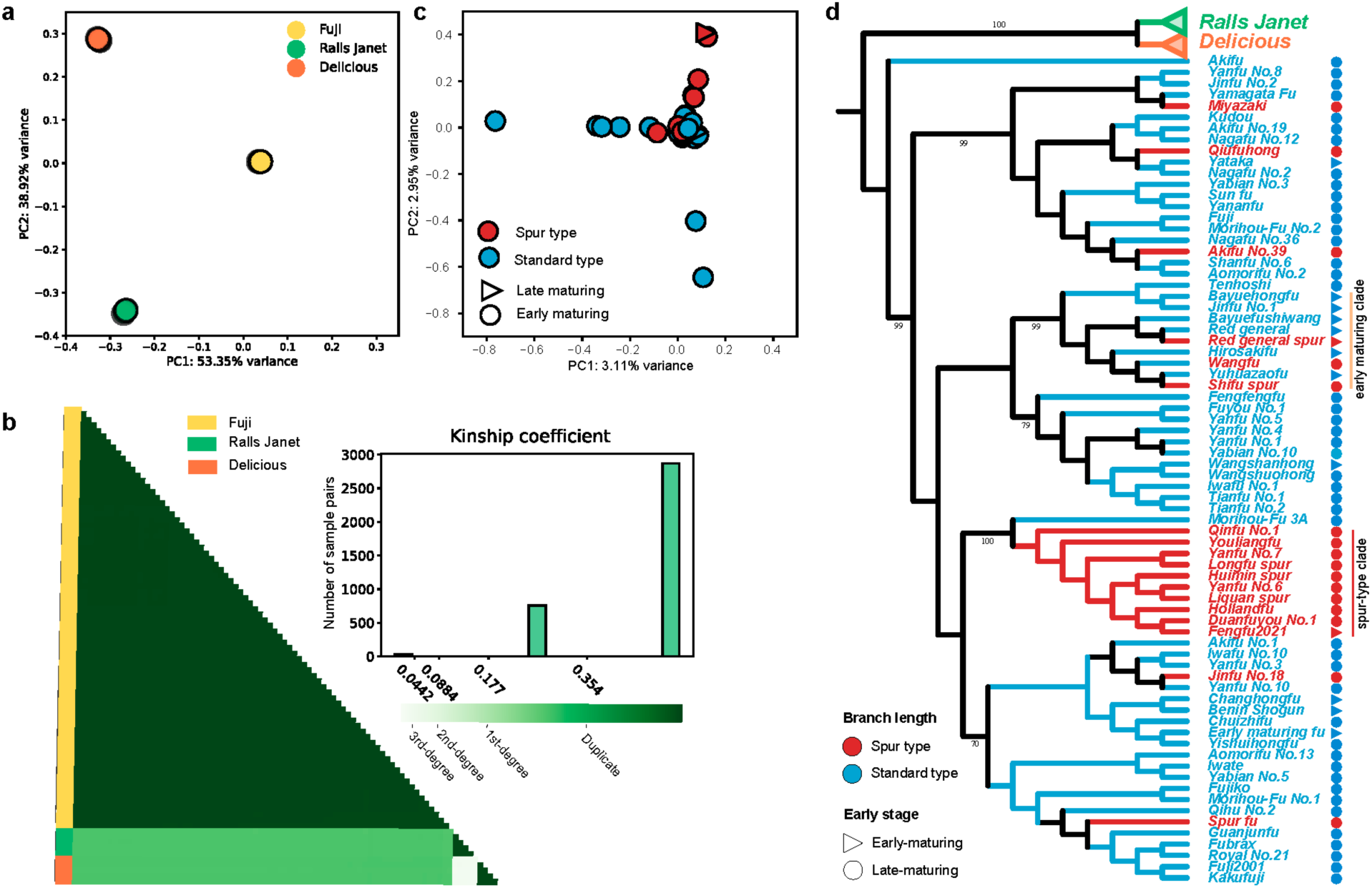
Population structure of ‘Fuji’ varieties. a. PCA of 74 ‘Fuji’ clonal varieties and five individuals from each of the two parents, ‘Ralls Janet’ and ‘Delicious’, using all identified SNPs. **b** Kinship of ‘Fuji’ clonal varieties, and ‘Ralls Janet’ and ‘Delicious’ individuals. The histogram in the upper right corner shows the frequency distribution of the coefficient of relatedness among all individuals. **c** PCA of ‘Fuji’ varieties based on somatic SNPs. **d** Phylogenetic relationships among the ‘Ralls Janet’, ‘Delicious’, and 74 ‘Fuji’ varieties based on somatic SNPs.

To further analyze the genetic relationships of the clonally propagated varieties of ‘Fuji’, we performed PCA on the 74 varieties using somatic SNPs, and the results did not show a clear genetic stratification (**Fig. 3c**). Furthermore, we performed phylogenetic analysis to track the evolutionary relationship among these varieties. The results revealed that the spur-type and early-maturing varieties were distributed among multiple clades (**Fig. 3d**), indicating that both traits had been formed independently in varieties belonging to different clades. Notably, we found one clade that consisted of only spur-type varieties (referred to as the spur-type clade), which suggested that these ten varieties might descend from a common spur-type ancestor. Similarly, there was also a clade enriched with early-maturing varieties (referred to as the early-maturing clade).

### Genetic basis of spur-type and early maturation traits

Spur-type growth habit is an essential economic trait in apple production. The trunk height and crown width of the spur-type clonal varieties were shorter than those of standard-type ‘Fuji’ varieties. Meanwhile, spur-type varieties have a higher spur rate than the standard-type ‘Fuji’ varieties (**Fig. 4a**). To further investigate the difference in characteristics between spur-type and standard-type varieties, the spur-type ‘Liquan spur’ and the standard-type ‘Yanfu No. 8’ were used as the representative varieties. Both the average shoot length and the average internode length of ‘Liquan spur’ were significantly shorter than those of ‘Yanfu No. 8’ (**Fig. 4b** and **Supplementary Fig. 6**). The contents of gibberellin A_3_ (GA_3_) and GA_4_ were lower in ‘Liquan spur’ than in ‘Yanfu No.8’ (**Fig. 4c**). In the paraffin section, the cell length was significantly shorter in ‘Liquan spur’ than in ‘Yanfu No. 8’, while the cell number in the same area was significantly increased in ‘Liquan spur’ (**Fig. 4d-f**). These results strongly suggest that cell length and density are critical factors in determining the internode length of apple trees.

**Fig. 4.**
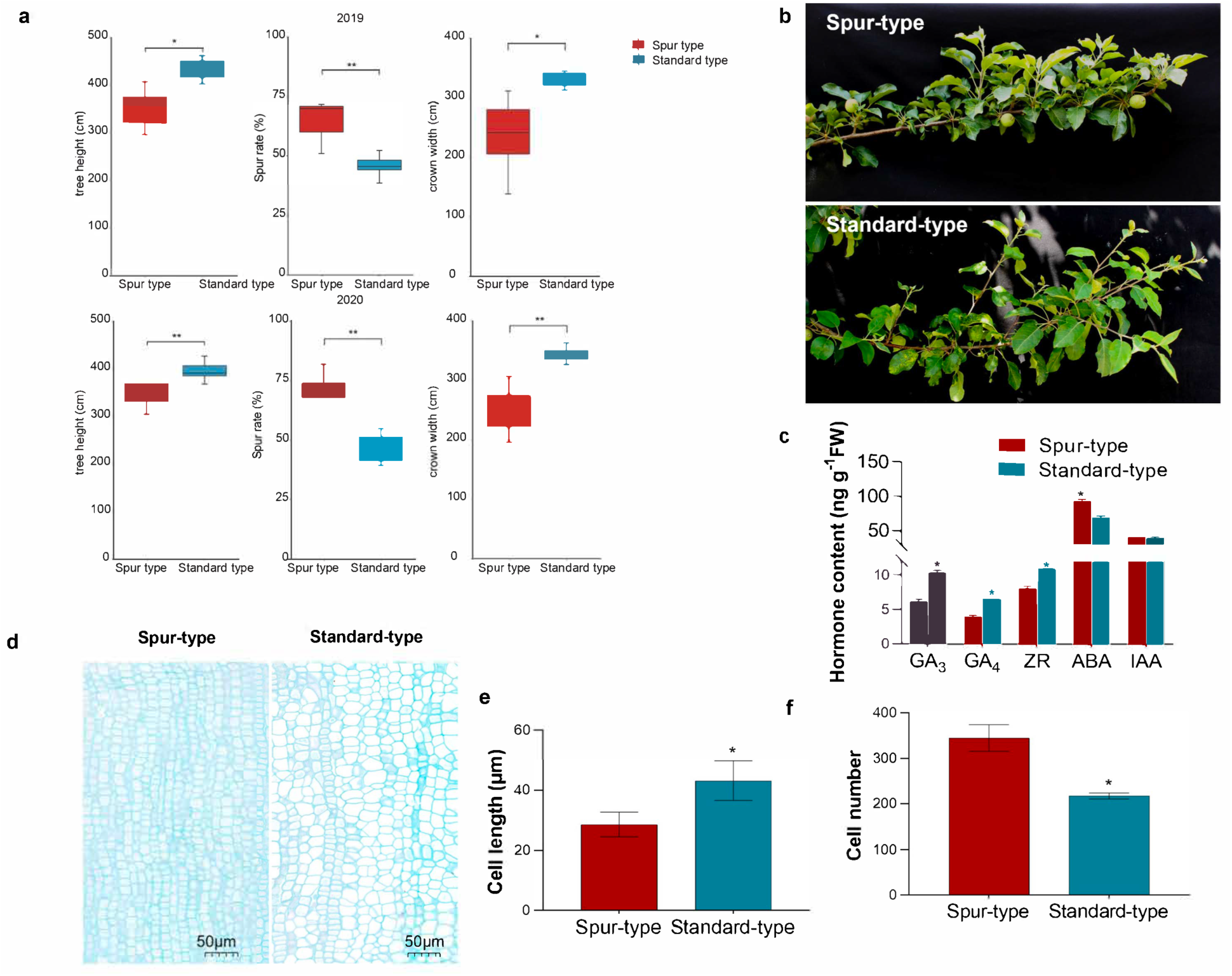
Physiological characteristics of standard and spur-type ‘Fuji’ varieties. a. Spur rate, crown width, and trunk height of standard and spur-type ‘Fuji’ varieties. Values represent the mean ± SE of three biological replicates, and * indicates a significant difference between means (*P* < 0.05). **b** Branches of standard-type ‘Yanfu No.8’ (Y8) and spur-type ‘Liquan spur’ (LQD). **c** Contents of GA_3_, GA_4_, trans-zeatin-riboside (ZR), abscisic acid (ABA), and indole-3-acetic acid (IAA) in Y8 and LQD. **d** Anatomical observations of a longitudinal section of the LQD and Y8 stems. **e,f** Cell length (**e**) and cell number (**f**) of the stem longitudinal sections of Y8 and LQD.

Phylogenetic analysis revealed two clades that were enriched with spur-type and early-maturing varieties, respectively (**Fig. 3d**). We detected clade-specific variants based on phylogenetic relationships to identify candidate somatic variants involved in spur-type and early maturation traits (**Supplementary Table 10**). We obtained 89 SNPs and eight indels specific to spur-type varieties in the spur-type clade, and 44 SNPs and two indels specific to the early-maturing varieties in the early-maturing clade (**Supplementary Tables 11 and 12**). We randomly selected 12 spur-type-specific SNPs and verified them in four spur-type and four standard-type varieties using Sanger sequencing (**Supplementary Table 13**). These clade-specific variants did not appear in other spur-type or early-maturing varieties, further supporting that the spur-type and early-maturing varieties that clustered into single clades, respectively, arose from a common ancestor. In contrast, spur-type and early-maturing varieties in other clades should have arisen independently.

### A TCP-like gene contributes to the spur-type morphology

We further examined whether somatic SVs contributed to the spur-type and early maturation phenotypes. By comparing the frequencies of each somatic SV in different clades, we did not identify any SVs specific to early-maturing varieties in the early-maturing clade. However, we found a 167-bp deletion specific to the varieties in the spur-type clade (**Fig. 5a**). This variant was present in all ten spur-type varieties in the clade but not in spur-type varieties in other clades or standard-type varieties. This 167-bp deletion overlapped with a 205-bp miniature inverted-repeat transposable element (MITE), encompassing 115 bp of the MITE sequence (**Fig. 5b**). The MITE existed as homozygous in other apple cultivars and their wild ancestor (**Supplementary Fig. 7**), indicating that the transposable element was inserted at an early stage. We selected five spur-type (from the spur-type clade) and five standard-type varieties for PCR validation for this deletion, and the verification rate was 100% (**Fig. 6a**). The SV was located in the promoter region of a TCP- like gene and contained a cis-acting element involved in GA response (**Fig. 5b**). It has been previously reported that genes from the TCP family are associated with traits such as internode length and branching^27,28^. Phylogenetic analysis showed that this TCP-like gene was highly homologous to Arabidopsis *TCP11* (**Fig. 5c**), so we named it *MdTCP11*.

**Fig. 5.**
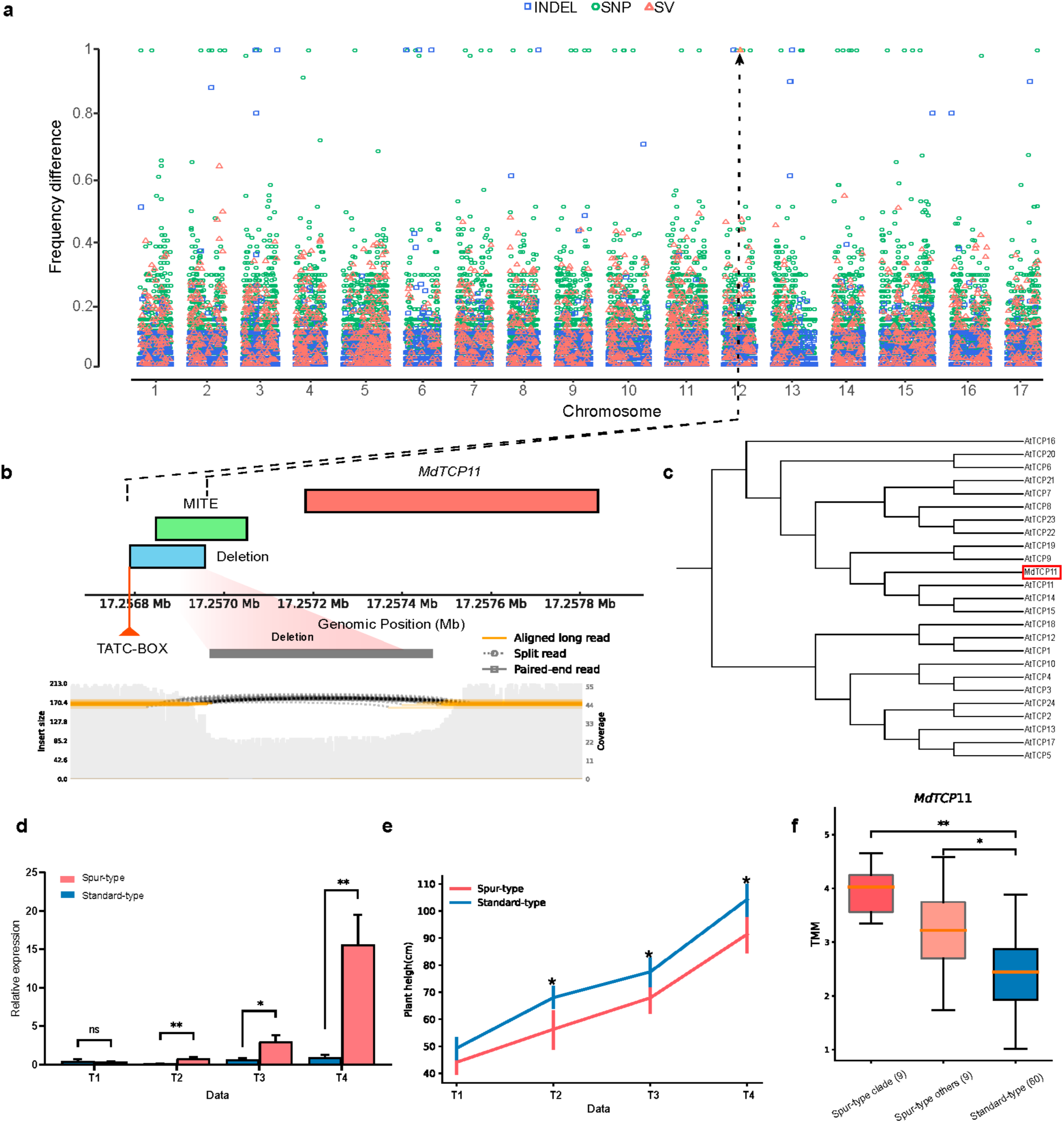
A 167-bp deletion and the expression of *MdTCP11* associated with spur-type growth habit. A. Frequency difference of somatic variants between the spur-type varieties in the spur-type clade and standard-type varieties. The y-axis represents the difference in frequency of samples with somatic mutations between the two groups. **b** Genomic positions of the 167-bp spur-type clade specific deletion, MITE and *MdTCP11*. The 167-bp deletion was located in the promoter of *MdTCP11* and contained a TATC-BOX. The read depth and sequence alignments around this deletion are shown below. **c** Phylogenetic analysis of *MdTCP11* and its homologs in Arabidopsis. **d** Expression of *MdTCP11* at different developmental stages of spur-type ‘Liquan spur’ and standard-type ‘Yanfu No.8’. T1, T2, T3, and T4: 90, 103, 112, and 134 days after germination, respectively. **e** Plant height of ‘Liquan spur’ and ‘Yanfu No.8’ at different stages. **f** Expression of *MdTCP11* in spur-type varieties in the spur-type clade, other spur-type and standard-type ‘Fuji’ varieties. * and ** indicate significant differences at P < 0.05 and 0.01, respectively (two-tailed Student’s t-test). ns, not significant.

**Fig. 6.**
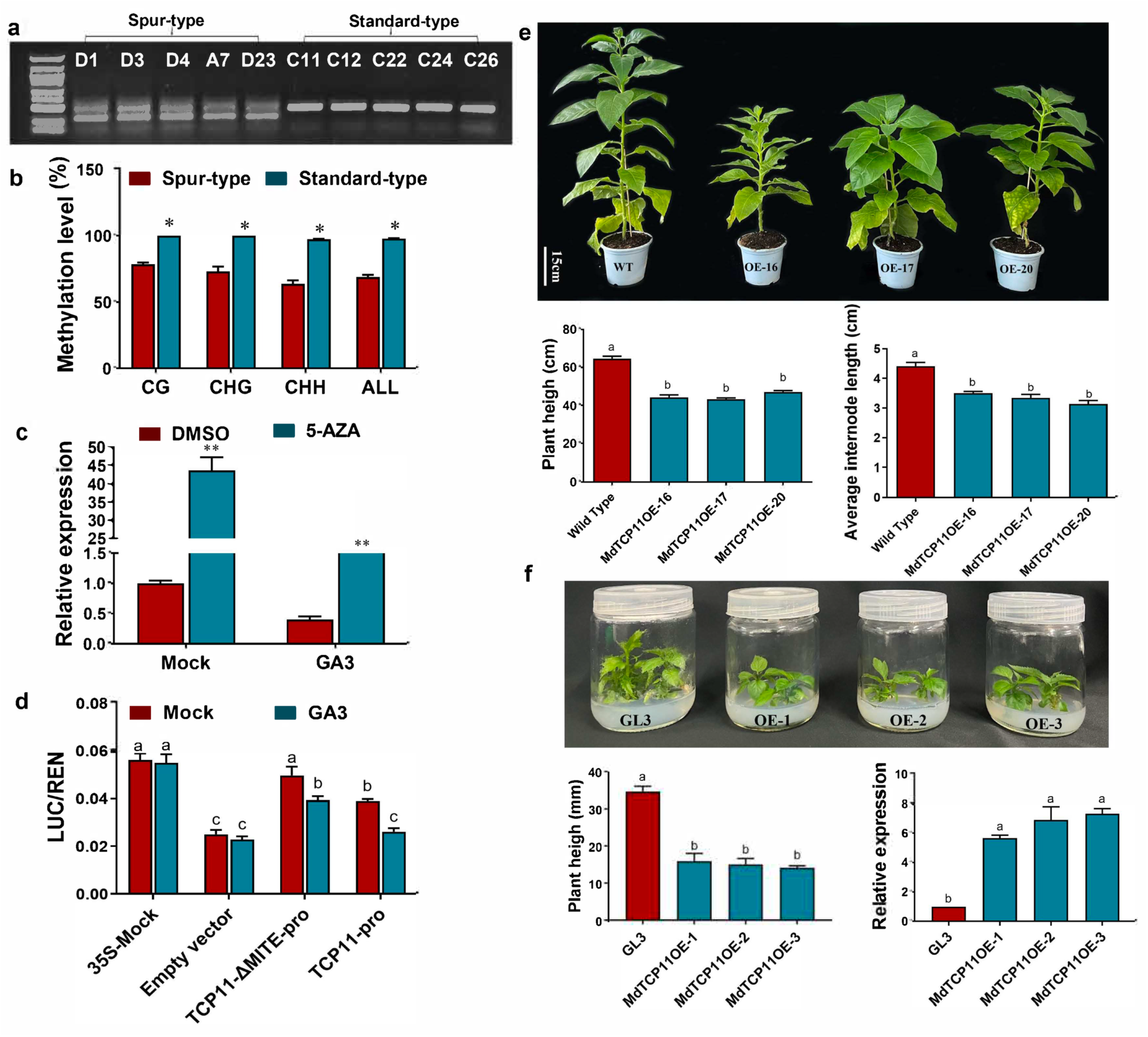
Promoter activity analysis and functional characterization of *MdTCP11*. a. PCR analysis of the 167-bp deletion in the *MdTCP11* promoter in different ‘Fuji’ clonal varieties. Detailed information of the varieties is provided in Supplementary Table 4. **b** Methylation levels of the MITE in standard-type and spur-type apple varieties. Data represent mean ± SE (n = 15). * and ** indicate significant differences at P < 0.05 and 0.01, respectively (Student’s t-tests). **c** Expression analysis of *MdTCP11* treated with 5-AZA under control and exogenous gibberellin conditions. DMSO, the solvent of 5-AZA, was used as the control. **d** LUC activity of the *MdTCP11* promoter treated with GA_3_. LUC-TCP11-ΔMITE-pro is the promoter of the spur-type variety. LUC-TCP11-pro is the promoter of the standard-type variety. Data represent mean ± SE (n = 6). **e** Morphology, plant height and internode length of transgenic *N. benthamiana* lines overexpressing *MdTCP11* and the wild type (WT). Data represent mean ± SE (n = 5). **f** Morphology of transgenic apple lines overexpressing *MdTCP11* and the wild-type ‘GL-3’. Means followed by different letters are significantly different at *P* < 0.05 based on two-way ANOVA.

Using time-series expression data of the typical spur-type variety ‘Liquan Spur’ and the standard-type variety ‘Yanfu No. 8’, we found that *MdTCP11* was differentially expressed between these two varieties at 103, 112 and 134 days after germination (**Fig. 5d**), consistent with the difference in plant height over time, indicating the potential role of *MdTCP11* in the formation of the spur-type morphology (**Fig. 5e**). RNA-Seq data further proved the differential expression of *MdTCP11* between the spur-type and the standard-type varieties (**Fig. 5f**). We also found that other spur-type varieties not in the spur-type clade also had higher expression of this gene compared with the standard-type varieties (**Fig. 5f**), although the 167-bp deletion was not detected in the promoter of *MdTCP11* in these spur-type varieties. The differential expression of *MdTCP11* was verified by qRT-PCR (**Supplementary Fig. 8**). We speculated that the upregulation of this gene in the spur-type varieties may contribute to the formation of the spur-type morphology.

To investigate the methylation status of the MITE, we performed site specific bisulfite sequencing (BS-seq) and found that the average DNA methylation level of the MITE in the standard-type varieties was higher than that in the spur-type varieties (**Fig. 6b**). To further determine the correlation between the MITE methylation and the expression level of *MdTCP11* with gibberellin treatment. We treated GL-3 plants with the DNA methylation inhibitor 5-AZA under control and exogenous gibberellin conditions. After treatment with 5-AZA, the expression of *MdTCP11* was significantly increased. Under exogenous gibberellin treatment, the expression of *MdTCP11* was increased by 5-AZA. These results further support the notion that MITE methylation is negatively correlated with *MdTCP11* expression under gibberellin treatment (**Fig. 6c**).

To further verify that the 167-bp deletion can respond to the GA treatment, we analyzed the promoter activity. The constructed vectors were transformed into *Agrobacterium* EHA105 and instantly transformed into apple calli. The infected apple calli were incubated on the GA3- containing medium, and then the LUC (luciferase) activities and GUS (β-glucuronidase) staining were assessed (**Fig. 6d** and **Supplementary Fig. 9**). The constitutive 35S promoter, *MdTCP1-ΔMITE-pro*, and *MdTCP11-pro* had vigorous GUS activities. After the GA_3_ treatment, GUS activity and LUC activity in *MdTCP1-ΔMITE-pro* and *MdTCP11-pro* were significantly inhibited. Furthermore, the GUS and LUC activities in *MdTCP1-ΔMITE-pro* were higher than those in *MdTCP11-pro.* Therefore, these results concluded that the expression activity of the *MdTCP1-ΔMITE-pro* promoter was higher than that of the *MdTCP11-pro* promoter, and that GA_3_ could inhibit the promoter activity of *MdTCP11*. To characterize the function of *MdTCP11*, we overexpressed *MdTCP11* in apple and *Nicotiana benthamiana*, which showed that overexpression of *MdTCP11* inhibited apple and *N. benthamiana* growth, and the plant height and average internode length were lower in *MdTCP11* overexpressing transgenic spple and *N. benthamiana* lines than in their corresponding wild types (**Fig. 6e,f**).

## Discussion

Somatic variations are prevalent in organisms, manifesting across all stages of growth and development. Their generation and accumulation have long been recognized as important factors in aging and the occurrence of tumors in humans. Therefore, studying the occurrence and development of somatic variations is of great significance. In plants, somatic variations are widely used in bud selection in fruit tree breeding^2^; however, the mechanisms underlying somatic variations remain largely unexplored.

The domestication of cultivated apple (*Malus domestica* Borkh.) has been driven by the hybridization of different wild ancestors^24^. Therefore, the apple genome has a high degree of heterozygosity, which challenges its genome assembly and downstream analyses. Reference genomes serve as the foundation for detecting genetic variants and are important resources for studying the genetic mechanisms underlying trait formation. The current apple reference genome is based on the cultivar ‘Golden Delicious’^16^, thus making it difficult to detect genetic variants in genomic sequences specific to ‘Fuji’. In addition, a single haploid genome has a bias toward the reference genome during read mapping, which interferes with subsequent analyses^29^, making it difficult to accurately identify causal variants. Trio-based methodologies employing HiFi sequencing currently represent the gold standard for achieving a fully phased genome assembly. This approach combines the high accuracy of trio-based phasing with the capability of long reads to bridge complex genomic regions, thereby offering unparalleled efficacy in segregating a genome into its paternal and maternal haplotypes^30^. In this study, we generated high-quality HiFi and ultra-long ONT reads, as well as Hi-C data for multiple ‘Fuji’ clonal varieties. Combined with short read data from the two parents of ‘Fuji’, we assembled a high-quality fully phased ‘Fuji’ genome, with individual haplotypes successfully assigned to the two parents of ‘Fuji’. This is in contrast with the recently published diploid ‘Fuji’ genome, in which the parental origins of individual haplotypes remain unresolved^31^ (**Supplementary Fig. 10**).

In plants, somatic variations can be passed on to the next generation. It has been shown that the proportion of somatic mutations passed on to the next generation in perennials (approximately 50%) is much higher than that of annual plants (approximately 10%). Therefore, somatic variations may have a greater contribution to the genetic diversity of perennials^25^. Due to the high heterozygosity of the ‘Fuji’ genome, the number of inherent germline variants present between the two haplotypes is much higher than the number of somatic variants, which could increase the chance of germline variants being wrongly identified as somatic variants. In addition, SVs between the two haplotypes can also complicate short-read alignments, resulting in false genetic variants. To systematically identify somatic variants in the ‘Fuji’ bud-sport varieties and study their functional impacts, we developed a pipeline based on the phased diploid genome inspired by the reference flow approach^32^. The pipeline aligns reads to the two haploid genomes and assigns them to the appropriate haplotypes according to the alignment quality. Compared with directly aligning reads to a single or diploid genome, this method obtains fewer mismatches, higher alignment scores, and thus higher alignment quality (**Supplementary Figs. 11-13**). This approach avoids most false positives caused by heterozygous germline variants inherited from Fuji’s common ancestor, allowing detection of somatic variants with higher confidence. Furthermore, we jointly used the long-read data from 13 varieties and the short-read data from 74 varieties to construct an SV dataset. This accurate and comprehensive collection of somatic variants provides an important foundation for identifying causal variants underlying interesting traits.

The bud sports of plants are usually caused by genetic and epigenetic variations. In contrast to epigenetic variations, genetic variations are stable and independent of environmental effects. In this study, the evolutionary relationships of ‘Fuji’ bud-sport varieties were reconstructed using high-confidence somatic SNPs. The results indicated that two important agronomic traits, spur-type growth habit and early maturation, had multiple independent formation events in different ‘Fuji’ bud-sport varieties. Interestingly, spur-type growth habit and early maturation each originated from one major event that affected multiple varieties. This is different from the result of cluster analysis using epigenetic variants, in which the spur-type and early-maturing varieties were scattered^33^. This result suggests two different strategies employed in the breeding of spur-type and early-maturing bud-sport varieties: 1) further selection from existing spur-type or early-maturing varieties to take advantage of the existing genetic variations, and 2) reliance on newly generated somatic variations from standard-type varieties to obtain spur-type or early-maturing traits.

Spur-type apples represent a germplasm resource with the characteristics of dwarfing, compactness, ease of management, early fruiting, high yield, etc., and are also an important germplasm that can be used for dwarf and dense apple cultivation. In apple, the ‘double dwarf’ cultivation method that combines spur-type apple varieties and dwarf rootstocks has become an important way for intensive dwarf and dense cultivation. Spur-type bud sports produce a dwarf tree architecture by reducing the duration of shoot vegetative growth, which is different from dwarfing rootstocks, which reduce the proportion of lateral buds that can develop into long shoots during the early development of trees^34,35^. Compared to standard-type varieties, spur-type varieties exhibited a reduced amount of annual extension growth, a tendency to produce spurs rather than shoots, and a smaller trunk cross-sectional area^36,37^. Consistent with these properties, we observed that spur rate and crown width significantly differed between standard-type and spur-type bud-sport varieties (**Supplementary Fig. 2**). Furthermore, the contents of GA_3_ and GA_4_ were lower in spur-type bud-sport varieties than in standard-type varieties (**Fig. 4c**). It has been reported that exogenous gibberellin application can promote shoot growth and increase internode length, and gibberellin biosynthesis inhibitors can inhibit internode growth^4^, indicating a positive correlation between endogenous hormone gibberellin and the internode length of branches in apple.

The formation of the spur-type trait is complicated, and spur-type varieties with different genetic origins should have different causal variants. In this study, we detected a number of genetic variants specific to the spur-type varieties that clustered together on the phylogenetic tree (**Supplementary Table 10**). These somatic variants may have contributed to the formation of this important trait. Among these somatic variants, we noticed one SV that was located in the promoter of a TCP family gene, *MdTCP11*. TCP transcription factors have been found to not only play a role in controlling cell proliferation and lateral organ development but also participate in plant hormone signaling and other processes. For example, overexpression of *MdTCP17* inhibits adventitious root formation in apple^38^, heterologous expression of *MdTCP12* inhibits axillary bud growth in Arabidopsis^39^, *TCP14* and *TCP15* affect internode length and leaf shape in Arabidopsis^28^, and Class I TCP-DELLA interactions at the inflorescence shoot apex determine the plant height^40^. Furthermore, *GrTCP11*, a TCP transcription factor in cotton, when expressed in *Arabidopsis thaliana* inhibits root hair elongation by downregulating genes in the jasmonic acid pathway^27^.

In this study, our transcriptome profiling data showed that *MdTCP11* was expressed at a higher level in spur-type varieties than in standard-type varieties. The promoter activity and methylation level analysis indicated that the expression of *MdTCP11* is negatively regulated by DNA methylation of the MITE in its promoter, and that the 167-bp deletion involving the MITE in the promoter could results in an increased expression level of *MdTCP11*. Notably, although the deletion was specific to the 10 spur-type varieties in the spur-type clade, we also observed that the expression level of *MdTCP11* was higher in other spur-type varieties than in standard-type varieties. This suggests that altered *MdTCP11* expression might also be involved in the emergence of other spur-type varieties, although the underlying genetic mechanism remains to be further investigated.

## Methods

### Plant materials and genome sequencing

A total of 84 apple samples, including 74 ‘Fuji’ clonal varieties, five ‘Ralls Janet’ varieties, and five ‘Delicious’ varieties (**Supplementary Table 4**), were collected at the Apple Demonstration Nursery of Yangling Modern Agriculture Technology Park (Northwest Agriculture & Forestry University), Shaanxi, China (34°52′N, 108°7′E). All the ‘Fuji’ varieties were grafted onto the M.26 rootstock in 2016. Genomic DNA was extracted from young leaves using the phenol-chloroform method. DNA libraries were constructed and sequenced on the Illumina NovoSeq platform.

PacBio SMRT libraries were prepared following the standard protocol provided by Pacific Biosciences (CA, USA) and sequenced on the PacBio Sequel II platform. Ultra-long Nanopore libraries were constructed with the SQK-LSK109 Ligation Sequencing 1D kit (Oxford Nanopore Technologies, UK), followed by the size-selection (>50 kb) via the SageHLS HMW system (Sage Science, USA), as per the manufacturer’s guidelines. The resulting libraries were sequenced on the PromethION platform. Genomic DNA of ‘Nagafu No.2’ was utilized to construct Hi-C libraries, which were then sequenced using the MGISEQ T7 platform.

### Genome assembly and annotation

The ‘Fuji’ genome was assembled using PacBio HiFi reads with the aid of ONT ultra-long reads, the parental short reads, and the Hi-C data. First, Hifiasm (v0.19.5)^30^ was applied independently to the HiFi reads from four ‘Fuji’ clonal varieties. During the assembly process, the parental short reads were integrated into the trio binning stage to generate haplotype-resolved contigs. Then the ONT ultra-long reads were integrated to further improve the continuity of assembly and produce the telomere-to-telomere level assembly. Next, Hi-C contact maps of the two haploid assemblies were generated using the 3D-DNA pipeline and Juicer tools^41^ with default parameters, which were used to help manual check and correction of potential misassemblies. Collinearity analysis between the ‘Fuji’ genome and the GDDH13 reference genome was performed using SyRI with default parameters^42^ and visualized using plotsr^43^. The 20-mers were collected from Illumina and HiFi reads of ‘Fuji’ and the two parents (‘Delicious’ and ‘Ralls Janet’) using Meryl (https://github.com/marbl/meryl). Subsequently, QV score and hamming error rates were calculated using Merqury^44^.

EDTA^45^ and RepeatModeler (http://www.repeatmasker.org/RepeatModeler/) were combined to annotate repeated sequences in the assembly. First, we used EDTA to identify LTR retrotransposons and DNA elements, and then RepeatModeler was used to generate *de novo* TE sequences. Finally, the two libraries were combined and imported into RepeatMasker (http://www.repeatmasker.org/RepeatMasker/) to scan the ‘Fuji’ genome assembly for repetitive elements.

Protein-coding genes were predicted from the ‘Fuji’ genome using the BRAKER3 pipeline^46^ that integrates RNA-Seq and protein homology information. RNA-Seq reads were aligned to the soft masked ‘Fuji’ genome using HISAT2 (v2.2.1)^47^. Protein sequences from *Malus domestica*, *Malus sieversii, Malus sylvestris, Pyrus communis* and *Prunus persica* were downloaded from GDR (https://www.rosaceae.org/) and aligned to the ‘Fuji’ genome as the protein homology evidence. As a complement to *de novo* gene predictions, we also mapped the gene annotations of GDDH13 onto the ‘Fuji’ genome using Liftoff (v1.6.3)^48^.

### Variant calling

Illumina paired-end reads were filtered and trimmed using fastp^49^ (v0.22.0). To detect germline variants, the cleaned reads were mapped to one haploid genome of ‘Fuji’ (Fuji_Ral) using BWA-MEM with default parameters^50^. The HaplotypeCaller module of GATK^51^ was utilized to call variants (SNPs and small indels). The resulting raw variants were then filtered using GATK with parameters ‘QD > 2.0 || MQ > 40.0 || FS > 60.0 || SOR > 3.0 || MQRankSum > −12.5 || ReadPosRankSum > −8.0’. Heterozygous alleles shared by more than 80% of ‘Fuji’ varieties were retained for the germline variant sets.

To detect somatic variants, the cleaned reads were aligned to the two haploid ‘Fuji’ genomes (Fuji_Ral and Fuji_Del), respectively, using BWA-MEM, which resulted in a pair of BAM files for each sample. According to the alignment score and the edit distance to the two haploid ‘Fuji’ genomes, we excluded alignments with lower alignment scores and higher edit distances in one of the paired BAM files. After that, we obtained a pair of cleaned BAM files for each sample, in which most reads were assigned to a proper haploid reference. Thus, alignments exhibiting the same alignment quality on both haploid genomes remained in both BAM files. We then used GATK HaplotypeCaller to detect variants from the cleaned BAM files with default parameters. Raw variants were filtered using GATK using the same parameters for filtering raw germline variants, and variants shared by more than 80% of ‘Fuji’ varieties were excluded to further reduce false positives. Since a subset of reads were retained in both BAM files, some genetic variants were reported in both haploid genomes. We further employed CrossMap^52^ to merge and eliminate redundancies in the genetic variants detected in the two haplomes, thereby establishing the final somatic variant collection. All the variants were annotated using SnpEff^53^. The SIFT4G protocol^54^ was used to identify deleterious variants.

### Structural variant calling

To identify high-confidence SVs, we used the strategy of combining long reads with short reads. We first used minimap2^55,56^ to align the HiFi and Nanopore reads to the ‘Fuji’ genome, and then used Sniffles (v2.2)^57^ to detect SVs and generate accurate SV boundaries. Next, we utilized GraphTyper (v2.7.5)^58^ to genotype SVs using Illumina short read data from the 74 ‘Fuji’ clonal varieties based on the SV dataset created by Sniffles. To identify potential hemizygous regions in the genome, we first compared Fuji_Ral and Fuji_Del to detect insertions and deletions between these two haploid genomes using svim-asm (v1.0.3)^59^ and Minigraph (v0.19)^60^. These SVs between the two haploid genomes were verified in the 74 clonal varieties using the results from GraphTyper. The reliable sequences with high repeatability (identified in more than 80% of samples) were preserved as the final diploid genome hemizygous sequence collection.

### Population genetic analyses

To infer the genetic relationships among our collected samples, we calculated the pairwise robust kinship estimator using the KING software^61^ with all SNPs. Principal component analysis (PCA) was performed using EIGENSOFT^62^. A phylogenetic tree was constructed using iqtree (v2.0.3)^63^ with the ‘MFP+ASC’ model and 1,000 bootstraps^64^, and then visualized using iTOL^65^.

### Identification of clade-specific variants

The ‘Fuji’ population was grouped according to their phenotypes and phylogenetic relationships (**Fig. 3d**). The frequency difference of each somatic variant between standard-type and spur-type varieties in the spur-type clade, as well as between late-maturing and early-maturing varieties in the early-maturing clade, were calculated. Variants specific to spur-type or early-maturing varieties were identified and then annotated using SnpEff (v5.0)^53^.

### Transcriptome and comparative genomic analysis of *MdTCP11*

Shoot tips from the ‘Fuji’ clonally propagated varieties were collected for RNA-Seq analysis. Total RNA was isolated using the Tiangen Total RNA Extraction Kit (DP441, Beijing, China), followed by mRNA enrichment employing magnetic beads conjugated with oligo(dT). Subsequently, RNA-Seq libraries were constructed using the NEBNext Ultra RNA Library Prep Kit and sequenced on the Illumina HiSeq 2500 platform. The raw RNA-Seq data were filtered using fastp (v0.22.0)^49^ with default parameters. The resulting cleaned reads were mapped to the Fuji_Ral genome using STAR (v2.7.4)^66^. Following read mapping, raw counts of each gene were derived using featureCounts^67^, and then normalized to TMM (Trimmed Mean of M-values) using the R package EdgeR^68^.

qRT-PCR analysis was performed using the CFX Connect Real-Time PCR Detection System (USA) and 2× SYBR Green Pro Taq HS Premix II (Accurate Biotechnology, Hunan, China), with three biological replicates and three technical replicates. Primers used for qRT-PCR are listed in **Supplementary Table 14**. Relative transcription levels were calculated according to the 2^−ΔΔCt^ method^69^.

To investigate the presence or absence of the MITE in the promoter of *MdTCP11* across various apple species, we retrieved genome assemblies and annotation files in gff3 format of 13 *Malus* accessions from NCBI (accession numbers PRJNA869488^14^, PRJNA927238^14^, and PRJNA591623^17^), and used the RepeatMasker to annotate the repeat sequences in these 13 apple genomes using the previously constructed TE library. *MdTCP11* orthologous gene pairs were identified using MCScan (Python version)^70^ with default parameters and collinearity relationships were then plotted based on the results of MCScan. Here, the 2-kb sequences upstream of the start codon were defined as promoter sequences. Finally, we integrated the results of gene structure annotation and repeat annotation to check whether MITE was present in the promoter of the *MdTCP11* gene among these 13 apple genomes.

### Morphological measurements and anatomical observations

Anatomical observations were performed using previously described protocols^71–74^. Stems of ‘Yanfu No. 8’ and ‘Liquan spur’ were fixed, dehydrated, and embedded in paraffin, as described in a previous study^74^. Sections were cut with a microtome (Leica DM2000, Germany) and stained with 0.1% toluidine blue (TB; Sakai 1973). The cell length and cell number of the stem longitudinal sections were measured, with three biological replicates from each line. All spur-type varieties were grafted onto the M.26 rootstock, and morphological measurements were conducted at the end of the growing season. The height and crown width of three trees of each variety were measured with a meter ruler and caliper. The number of 1-year-old spurs (< 5 cm) graving on the 2-year-old branches for each variety was counted, and the spur rate was calculated as the number of spur shoots divided by the number of total shoots. Three randomly selected main branches of each tree were used to count the spur rate. ‘Liquan spur’ and ‘Yanfu No. 8’ were used to represent spur-type and standard-type varieties, respectively. Three trees of each of these two varieties were used to measure the length and internode length of 1-year-old shoots on three randomly selected main branches of each tree. The average internode length was calculated by dividing the overall shoot length by the number of internodes.

### Plant material and growth condition

Tissue-cultured GL-3 (the progeny of ‘Royal Gala’ apple, which was used as the background for apple transformation) and transgenic plants were subcultured every 4 weeks on Murashige & Skoog (MS) medium (4.43 g/L MS salts,30 g/L sucrose, 0.2 mg/L 6-BA,0.2 mg/L IBA, and 7.5 g/L agar, pH 5.8) under long-day conditions (14 h light :10 h dark) at 25°C. Apple calli (*Malus domestica* cv. ‘Orin’) were grown on MS medium containing 0.4 mg/L 6-BA and 1.5 mg/L 2,4-D at 25 °C under dark conditions and were subcultured every 2 week.

### Hormone extraction and measurement

Shoot tips of ‘Yanfu No. 8’ and ‘Liquan spur’ were collected at 60 days after flowering. Hormones GA_3_, GA_4_, ZR, ABA, and IAA were purified and extracted from the collected samples following a previously described procedure^22^. Three biological replicates were used for each sample. The detection and analysis of these hormones were performed with high-performance liquid chromatography (Waters 2498/UV; Visible Detector, Shaanxi, China)^23^. External standards (GA_3_, GA_4_, ZR, ABA, and IAA) were purchased from Sigma (USA) and used for quantitative analyses.

### GUS staining

The promoter of *MdTCP11* (*MdTCP11-pro*) and the *MdTCP1-ΔMITE-pro* sequence were inserted into *pCAMBIA1381-GUS*, and 35S:GUS and the empty *pCAMBIA1381-GUS* vector were used as the positive and negative controls, respectively. The resulting constructs were transferred into *Agrobacterium tumefaciens* (strain EHA105) using *Agrobacterium*-mediated transformation to obtain an instantaneous transgenic apple callus. GUS staining was performed after 3 days of dark culture using 5-bromo-4-chloro-3-indolyl-β-glucuronide (X-gluc) as a substrate, as previously described^75^. The apple calli used for injection were selected from the tissue culture for two weeks.

### Dual-luciferase assays

The *MdTCP11-pro* and *MdTCP1-ΔMITE-pro* sequences were inserted into the pGreen II 0800-LUC vector, with the 35S promoter serving as a positive control and the empty pGreen II 0800-LUC vector as a negative control. The vectors were transformed into *Agrobacterium* strain EHA105 and then transformed into apple calli. After 3 days of culture in the dark, LUC and REN activities were quantified using a dual-luciferase reporter assay system (Promega, E1910). At least ten biological replicates were performed for each transformation, and the ratios of LUC to REN were calculated for treatments and controls to assess the activity of *MdTCP11*.

### Subcellular localization

The CDS of *MdTCP11* was inserted into the pC2300-GFP vector, which was transformed into *Agrobacterium* strain GV3101, and then transformed into *N. benthamiana*. After 3 days of culture in the dark, GFP fluorescence was detected using a confocal laser-scanning microscope with excitation at 488 nm (Zeiss LSM 510 Meta, Jena, Germany).

### DNA methylation assay

For locus-specific BS-seq, DNA was extracted from ‘Nagafu No.2’ (standard-type) and ‘Yanfu No.6’ (spur-type) leaves using a Super Plant Genomic DNA Kit (TIANGEN, DP360, China). Approximately 300 ng of DNA was treated with bisulfite using an EZ DNA Methylation-Gold Kit (Zymo Research, D5005, USA) and amplified by PCR using Takara Premix Ex TaqTM Hot Start Version (Takara Research, RR030A, Japan). The amplified products were recovered from an agarose gel using a Universal DNA Purification Kit (TIANGEN, DP214, China). The purified product was cloned into the pBM16A vector using a pBM16A Topsmart Cloning Kit (BIOMED, China). At least 15 clones were sequenced per genotype using the Sanger sequencing technology. The final results were analyzed using the Kismeth online software^76^. 5-AZA (Sigma, A3656, USA) treatment was performed according to the previous description and slightly modified^77^. The subcultured GL-3 were transferred to MS medium containing 7ug/L of 5-AZA or DMSO for 3 weeks under long-day conditions (14 h light, 10 h darkness) at 25 ℃, followed by treatment with 2mg/L of GA_3_ for 24h.

### Genetic transformation

The *pC2300-MdTCP11-GFP* vector was transformed into *Agrobacterium* strain EHA105, and then transformed into apple and *Nicotiana benthamiana* leaves. Transgenic apple and *N. benthamiana* microshoots were maintained in *in vitro,* and subcultures were performed every 4 weeks. Apple and *N. benthamiana* were grown in a growth room under long-day conditions (16 h light/8 h dark, 25 °C). Plant height and internode length were measured in transgenic *N. benthamiana* lines overexpressing *MdTCP11* and wild type (WT), and plant height was measured in transgenic apple lines overexpressing *MdTCP11* and WT (‘GL3’). Five biological replicates from each line were conducted.

## Data availability

Raw sequencing data and genome assemblies have been deposited in China National Center for Bioinformation (CNCB) under the BioProject accession number PRJCA024962.

## Acknowledgments

We thank Dr. L. Wang from Huazhong Agricultural University for help with the genomic analyses, Prof. L. Wang from Nanjing University for the critical reading of the manuscript, Prof. C.X. You from Shandong Agricultural University for providing the ‘Orin’ apple calli, Prof. Z.H. Zhang from Shenyang Agricultural University for providing tissue-cultured ‘GL-3’ plants and the high - performance computing platform of Northwest A&F University (HPC of NWAFU). This work was supported by grants from the National Natural Science Foundation of China (32372657 and 32372675), the National Key Research and Development Project (2023YFD2301002), Xinjiang Corps Science and Technology Program (2023AB077), the Cyrus Tang Foundation, and the China Apple Research System (CARS-27).

## Author contributions

D.Z., Y.J. and Z.F. designed and coordinated the study. Y.C., Yu Liu, L.T., and W.K. performed genomic and transcriptomic analyses. Y.C. and J.Mao performed the data analysis. X.G., Yu Liu, L.T., C.C., Yandong Liu, X.Z., C.C., S.L., X.S. and D.M. performed the experiments and data analysis. Z.Y., Z.L., F.M., Z.Z., W.L. and Q.L. provided samples. Y.C., J.Mao, D.Z., Z.F. and Yu Liu wrote the paper with contributions from T.F., J.Y., A.C., X.C., M.M.T., J.Ma, L.X., N.A., Y. L., C.Z., Y.H. and H. L.

