## Supplementary Figures for "Genome sequencing of ‘Fuji’ apple clonal varieties reveals evolutionary history and genetic mechanism of the spur-type morphology"

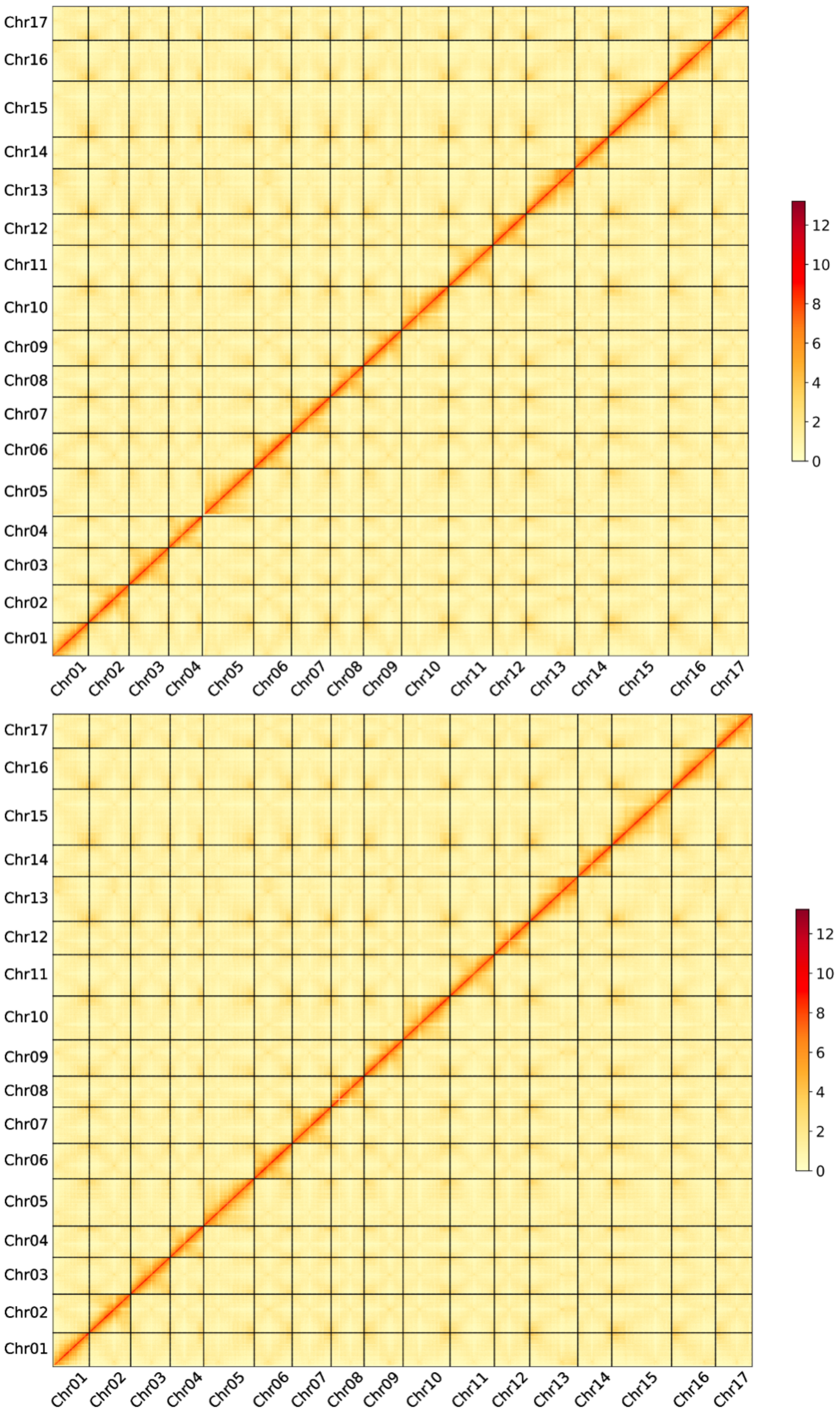


**Supplementary Figure 1.** Hi-C contact map of Fuji_Ral (top) and Fuji_Del (bottom) assemblies. The intensity of interactions was calculated using a bin size of 600 kb.

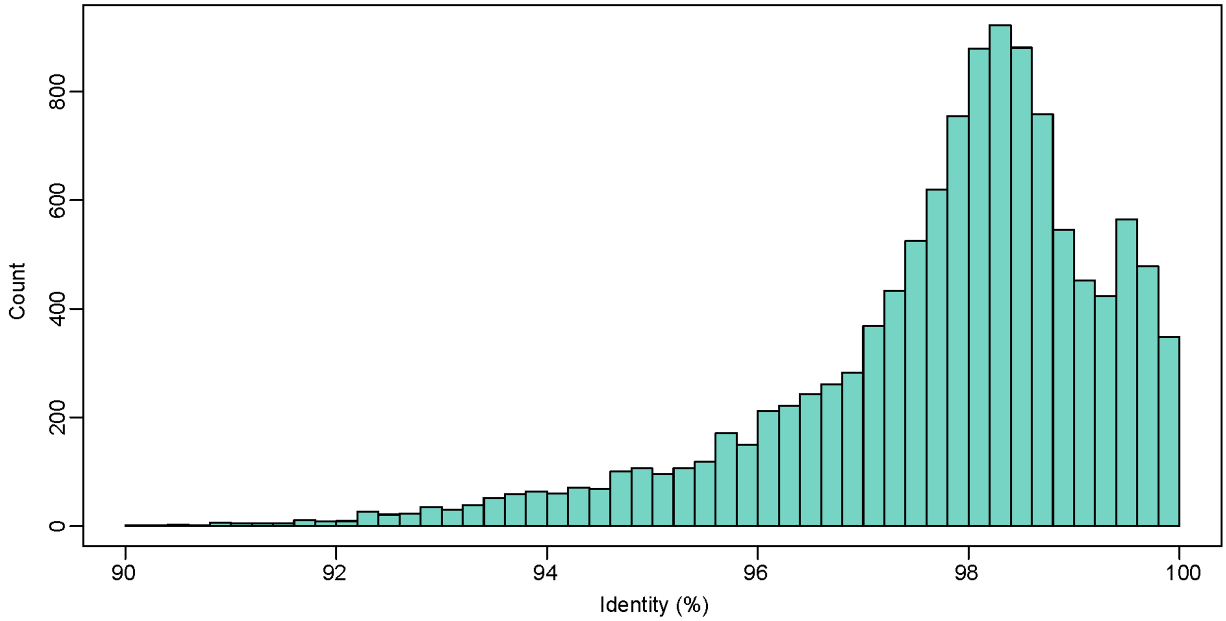


**Supplementary Figure 2.** Distribution of sequence identities in the alignment blocks between the two haploid genomes of ‘Fuji’. The x-axis represents sequence identity, and the y-axis indicates the number of windows with the corresponding sequence identity.

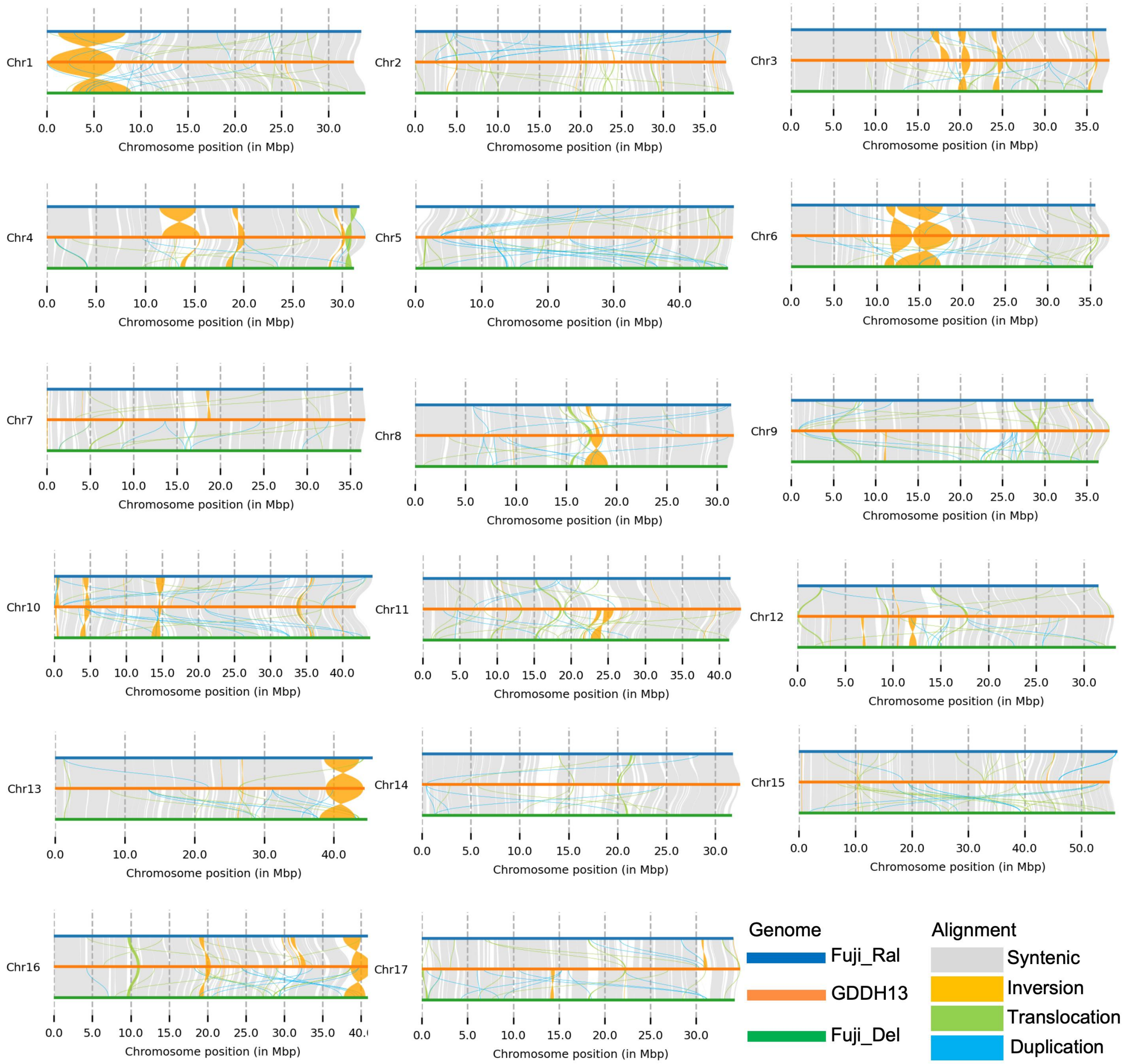


**Supplementary Figure 3.** Collinearity between the 'Fuji' genome and the apple reference genome (GDDH13).

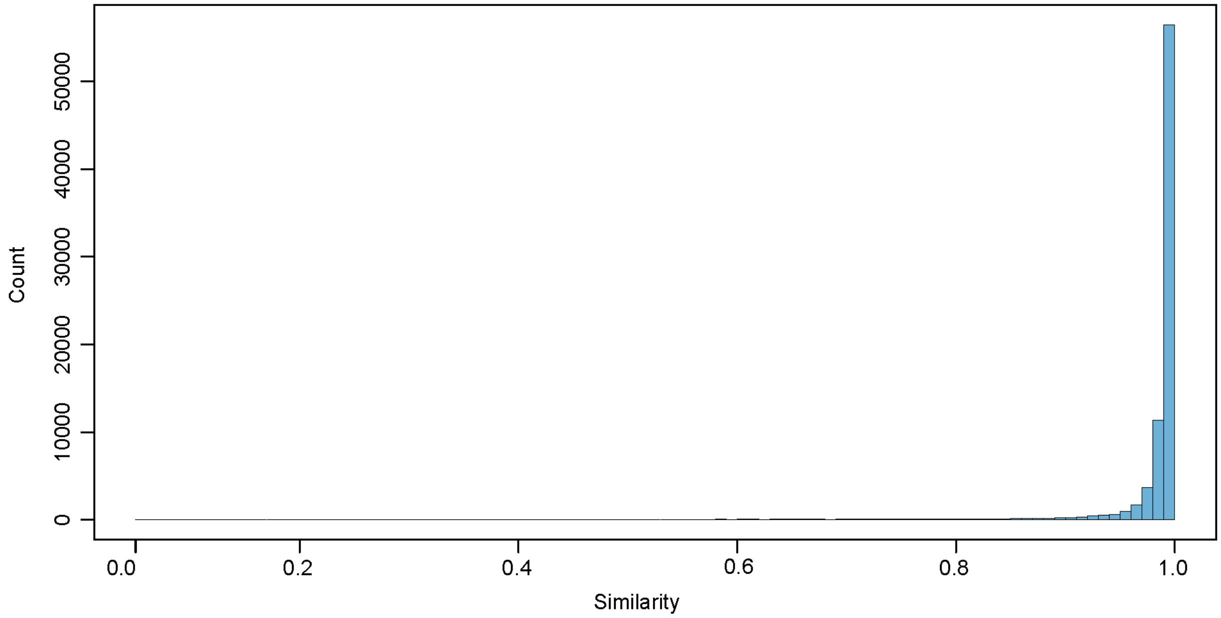


**Supplementary Figure 4.** Distribution of sequence identities of protein-coding genes between Fuji and GDDH13.

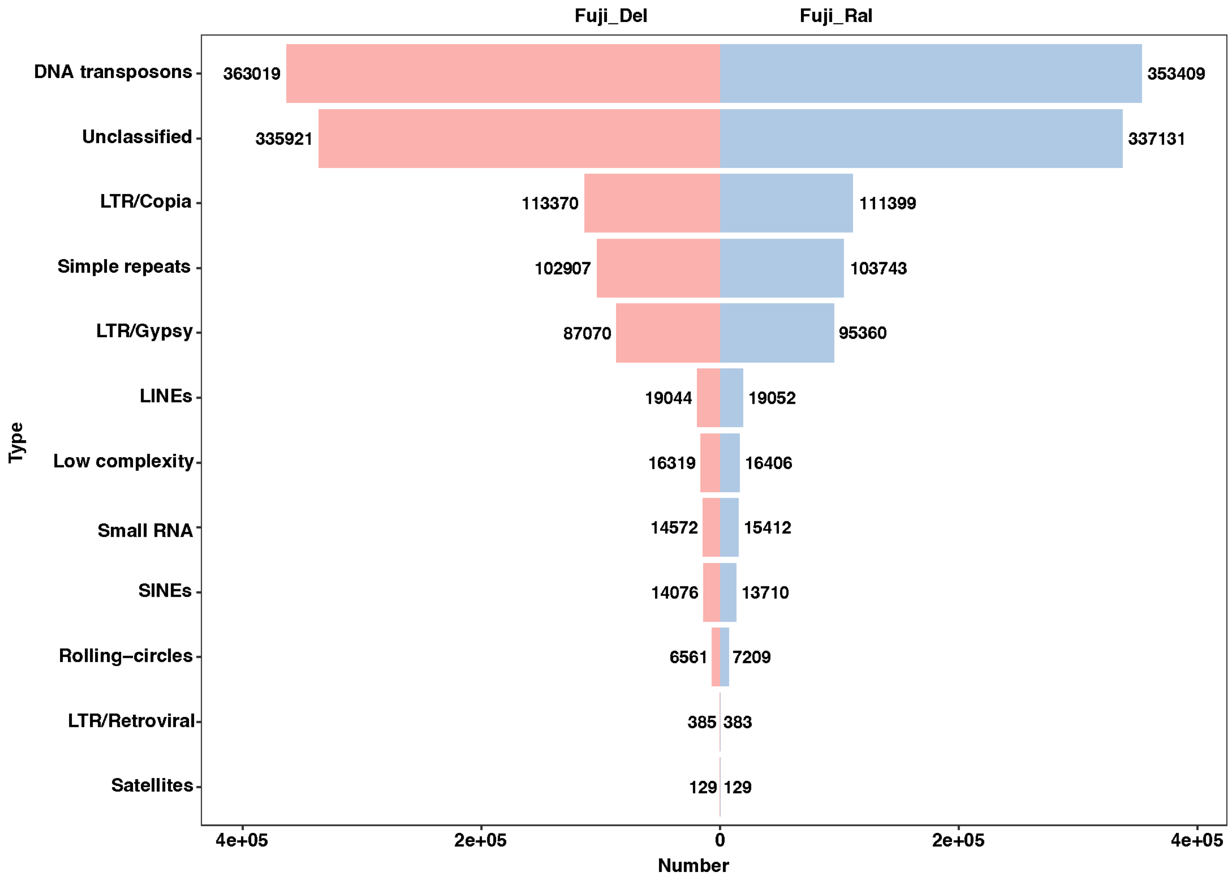


**Supplementary Figure 5.** Type and number of repeat sequences in the two 'Fuji' haploid genomes.

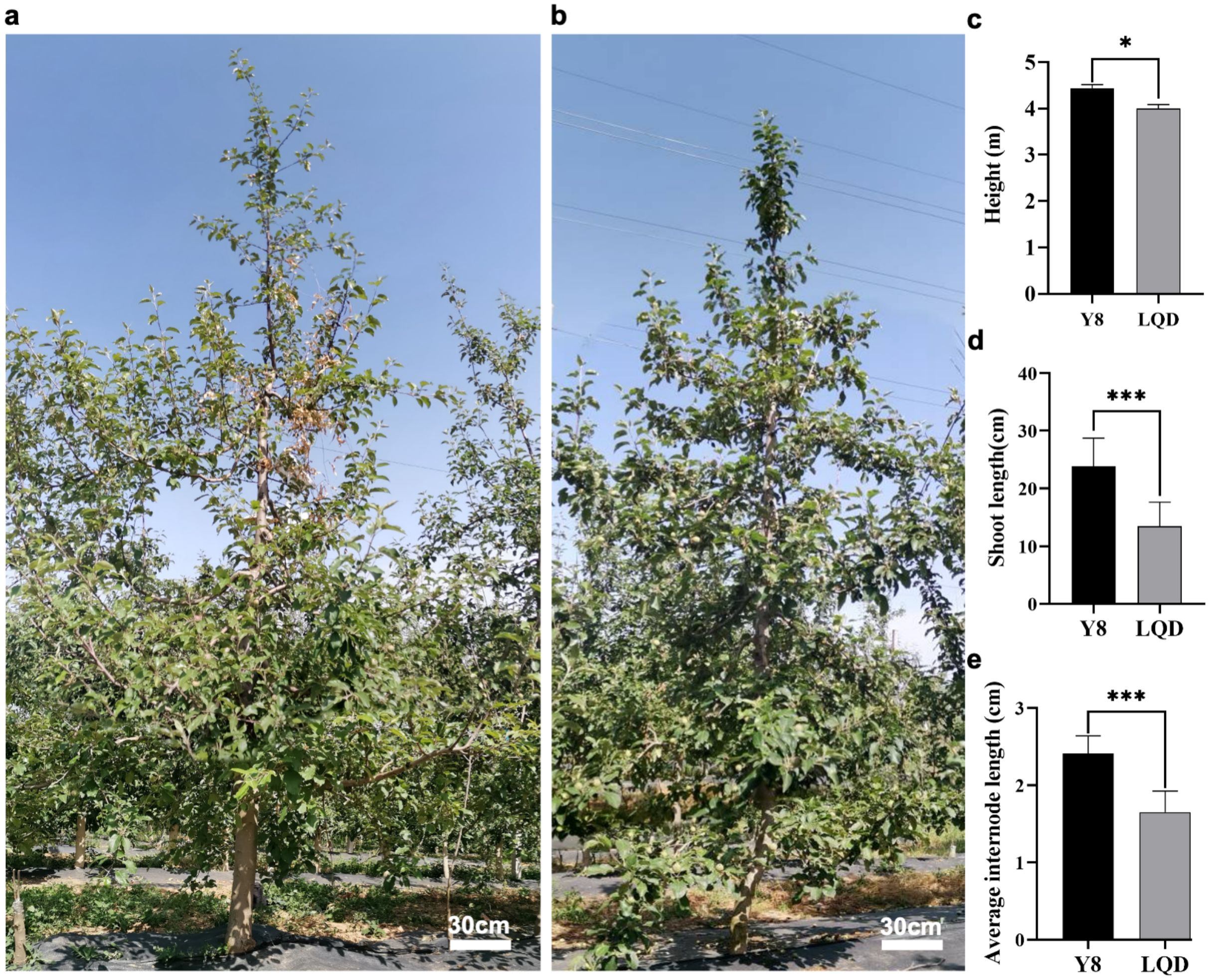


**Supplementary Figure 6.** Phenotype characterizations of standard and spur-type apple varieties**.** **a,b** Plants of standard (**a**) and spur-type (**b**) apple varieties. **c-e** Height (**c**), shoot length (**d**), and internode length (**e**) of standard-type variety ‘Yanfu No.8’ (Y8) and spur-type variety ‘Liquan spur’ (LDQ). Values represent mean ± SE of three biological replicates. * and *** indicate significant difference between means (*P* < 0.05 and *P* < 0.001, respectively).

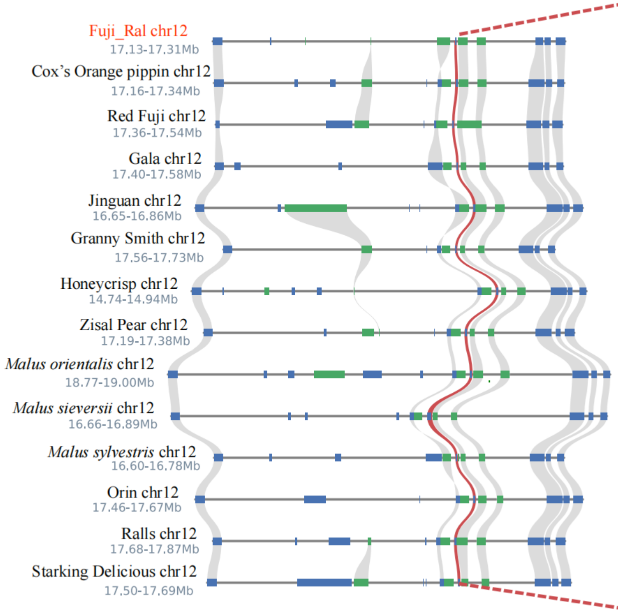


| Species | Gene | TE in promoter region |
|---|---|---|
| Fuji | MdTCP11 | √ |
| Cox's Orange Pippin | COP012G0103200 | √ |
| Red Fuji | FJ012G0099300 | √ |
| Gala | Mdg_12g011190 | √ |
| Jiguan | JG012G0098600 | √ |
| Granny Smith | GS012G0106900 | √ |
| Honeycrisp | HC012G0090500 | √ |
| Zisai Pear | ZP012G0107700 | √ |
| *Malus orientalis* | MO012G0121500 | √ |
| *Malus sieversii* | Msi_12g010660 | √ |
| *Malus sylvestris* | Msy_12g010770 | √ |
| Orin | OR012G0103500 | √ |
| Ralls | RA012G0101200 | √ |
| Starking Delicious | SD012G0101500 | √ |

**Supplementary Figure 7.** Microsynteny of the *MdTCP11* genome regions among the 'Fuji' genome and genomes of other 13 apple accessions. *MdTCP11* and syntenic gene pairs are depicted in red, while the syntenic flanking genes are connected by gray lines. The table on the right lists the presence of the MITE in the promoter of *MdTCP11* across various genomes.

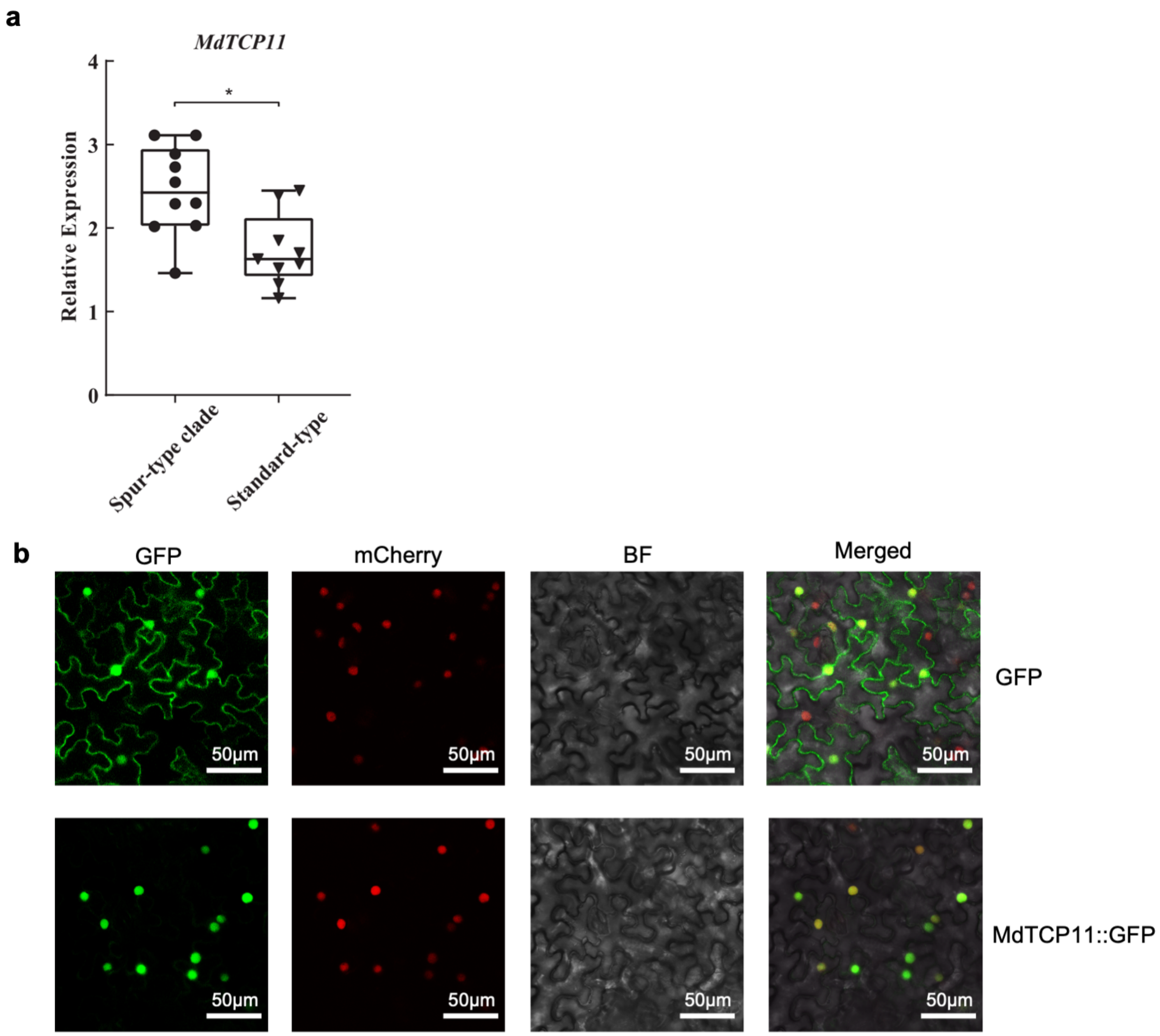


**Supplementary Figure 8.** Expression and subcellular localization of MdTCP11. **a** Expression of *MdTCP11*. **b** Subcellular localization of MdTCP11.

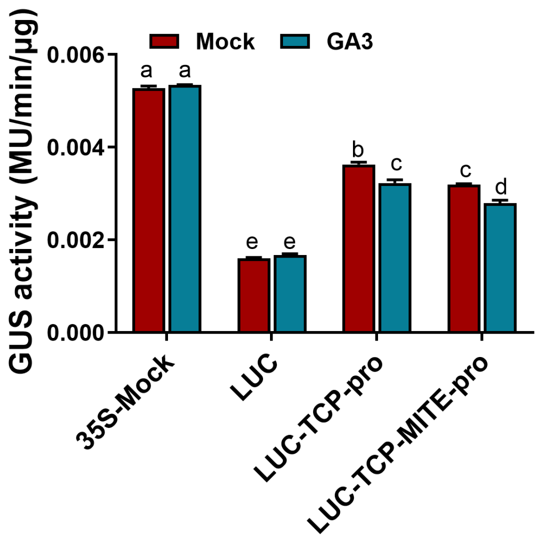


**Supplementary Figure 9.** Promoter activities of *MdTCP11* treated with $GA_3$. TCP11-ΔMITE-pro is the promoter of spur-type apple. TCP11-pro is the promoter of standard-type apple. Data represent mean ± SE (n = 6). Different letters above bar graphs indicate significant difference at $P < 0.05$.

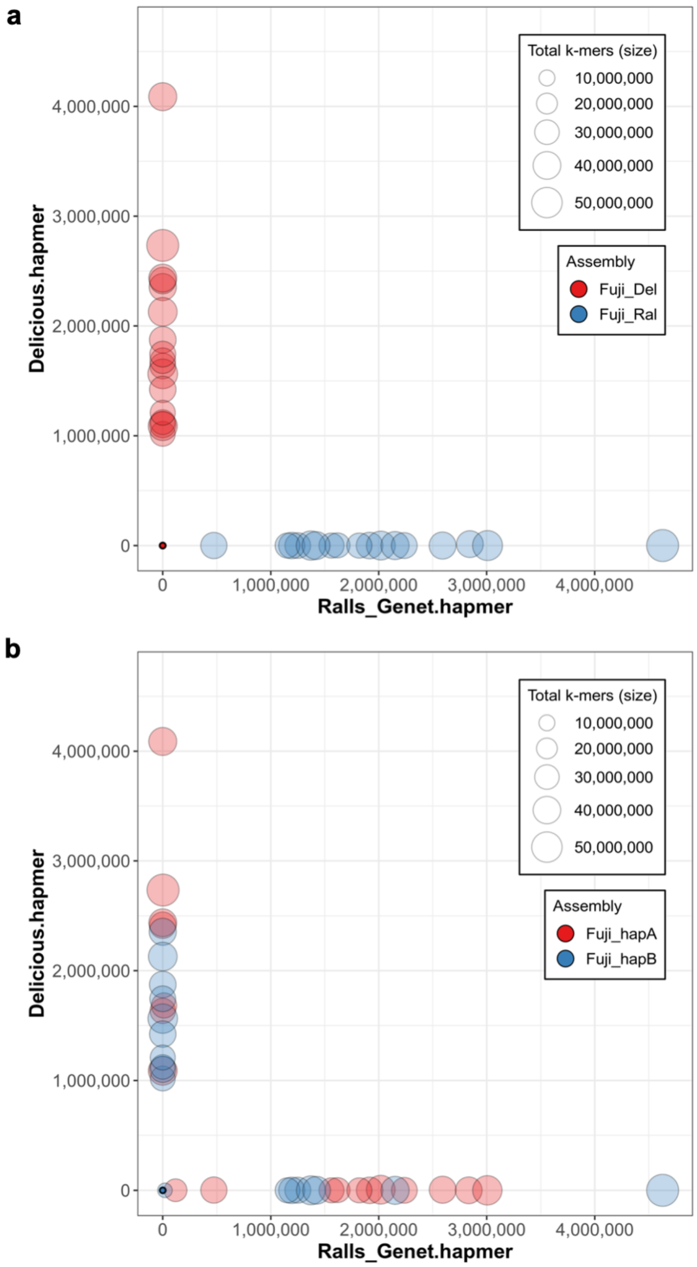


**Supplementary Figure 10.** Haplotype-specific k-mer **(**hap-mer) blob plots of Fuji assemblies. **a,b** Hap-mer blob plots of the assemblies Fuji_Ral and Fuji_Del (**a**) and previously published Fuji_hapA and Fuji_hapB assemblies (**b**). The chromosomes of the two haplomes are colored by red and blue, respectively. The x and y axes indicate the number of 'Ralls Genet' hap-mers and 'Delicious' hap-mers, respectively, that the chromosomes contain. The blob size is proportional to the total k-mers of each scaffold. This result indicates that Fuji_Ral and Fuji_Del are fully phased in our assembly, while the genetic origins are switched in some chromosomes of the previously reported Fuji_hapA and Fuji_hapB assemblies.

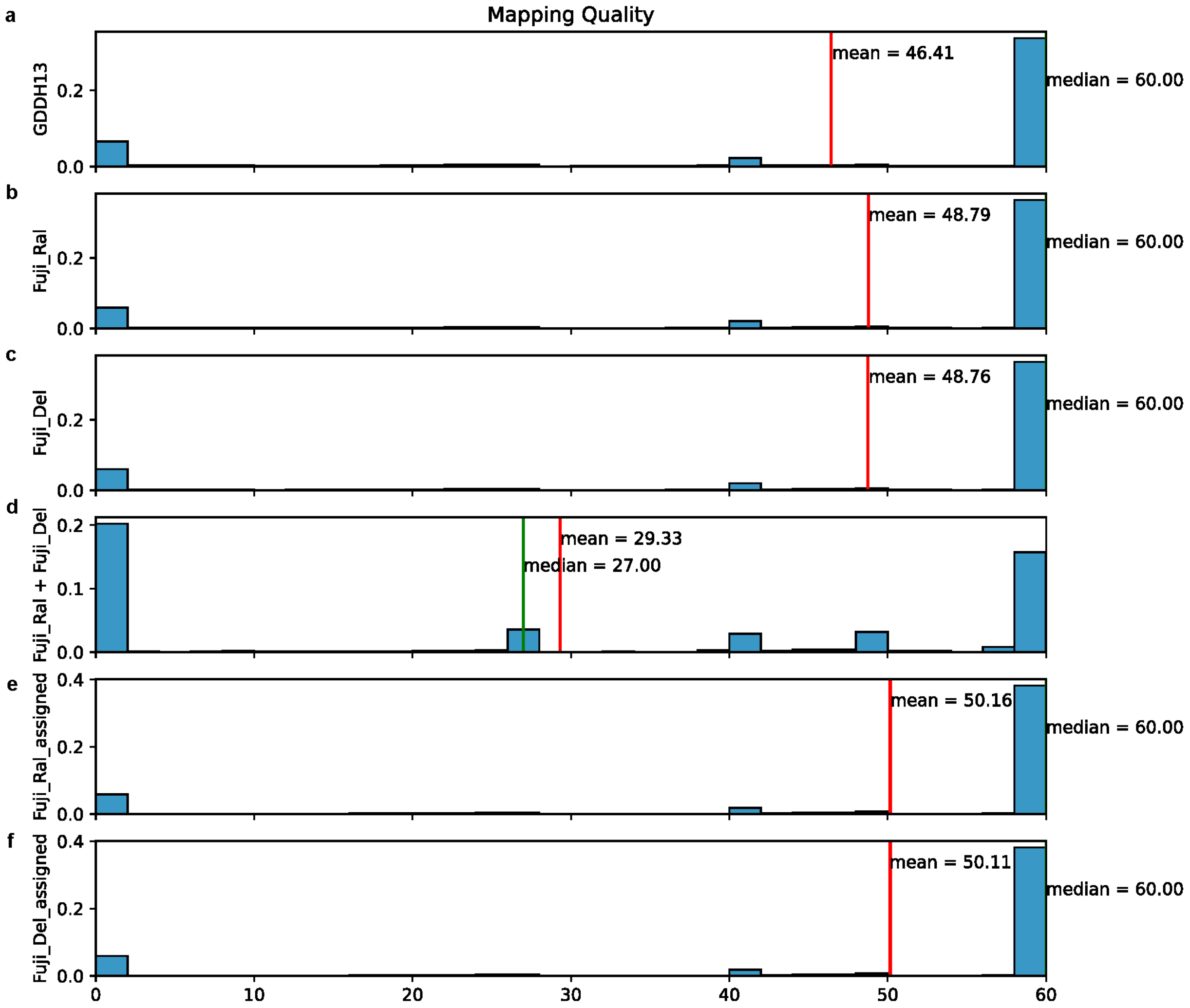


**Supplementary Figure 11.** Mapping quality using different references for read mapping. **a** Using GDDH13 as the reference genome. **b** Using Fuji_Ral as the reference genome. **c** Using Fuji_Del as the reference genome. **d** Using Fuji_Ral + Fuji_Del as the reference genomes without secondary assignments of reads. **e,f** Assigning reads to the appropriate haplotypes using the pipeline developed in this study.

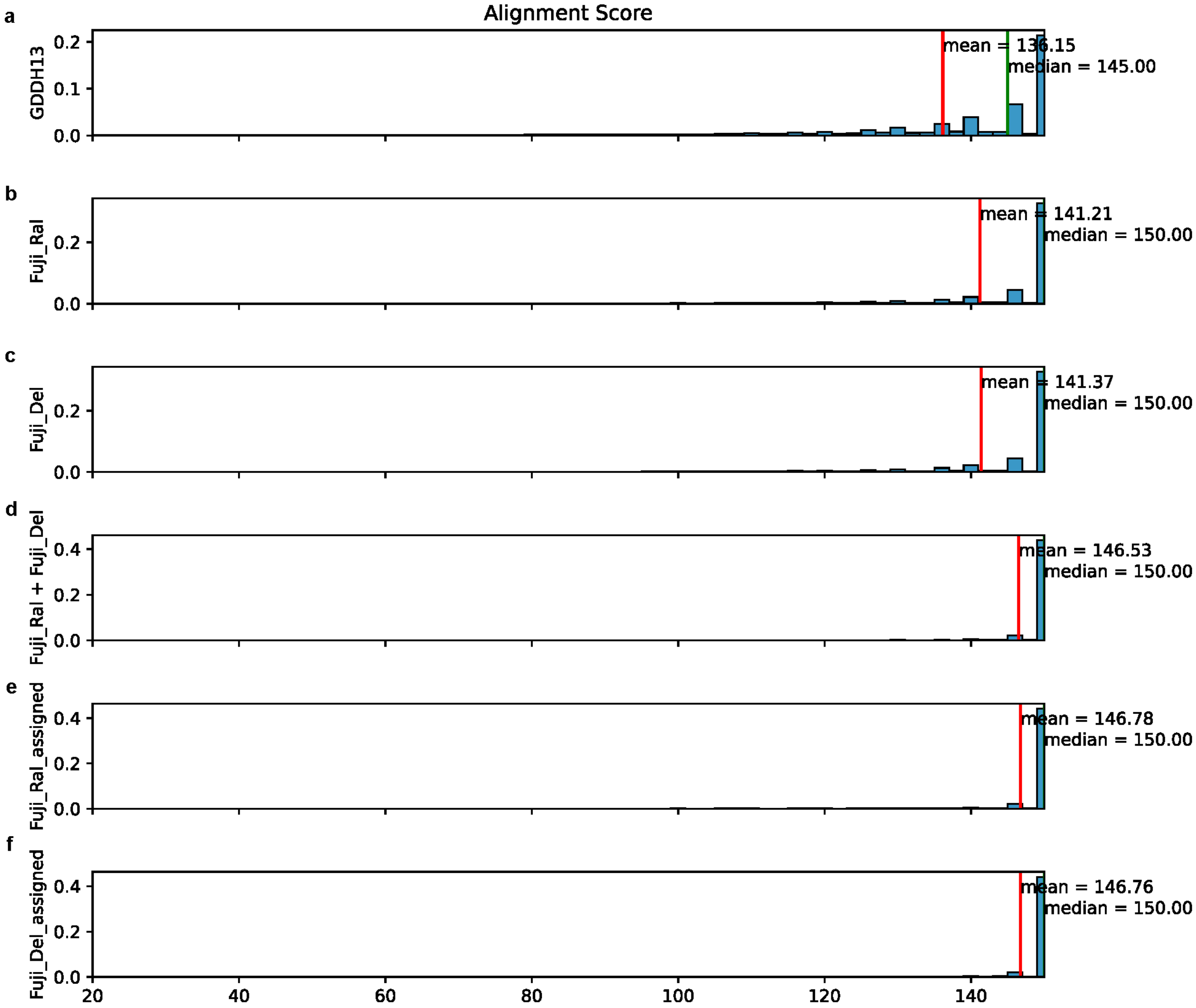


**Figure S12.** Alignment score using different references for read mapping. **a** Using GDDH13 as the reference genome. **b** Using Fuji_Ral as the reference genome. **c** Using Fuji_Del as the reference genome. **d** Using Fuji_Ral + Fuji_Del as the reference genomes without secondary assignments of reads. **e,f** Assigning reads to the appropriate haplotypes using the pipeline developed in this study.

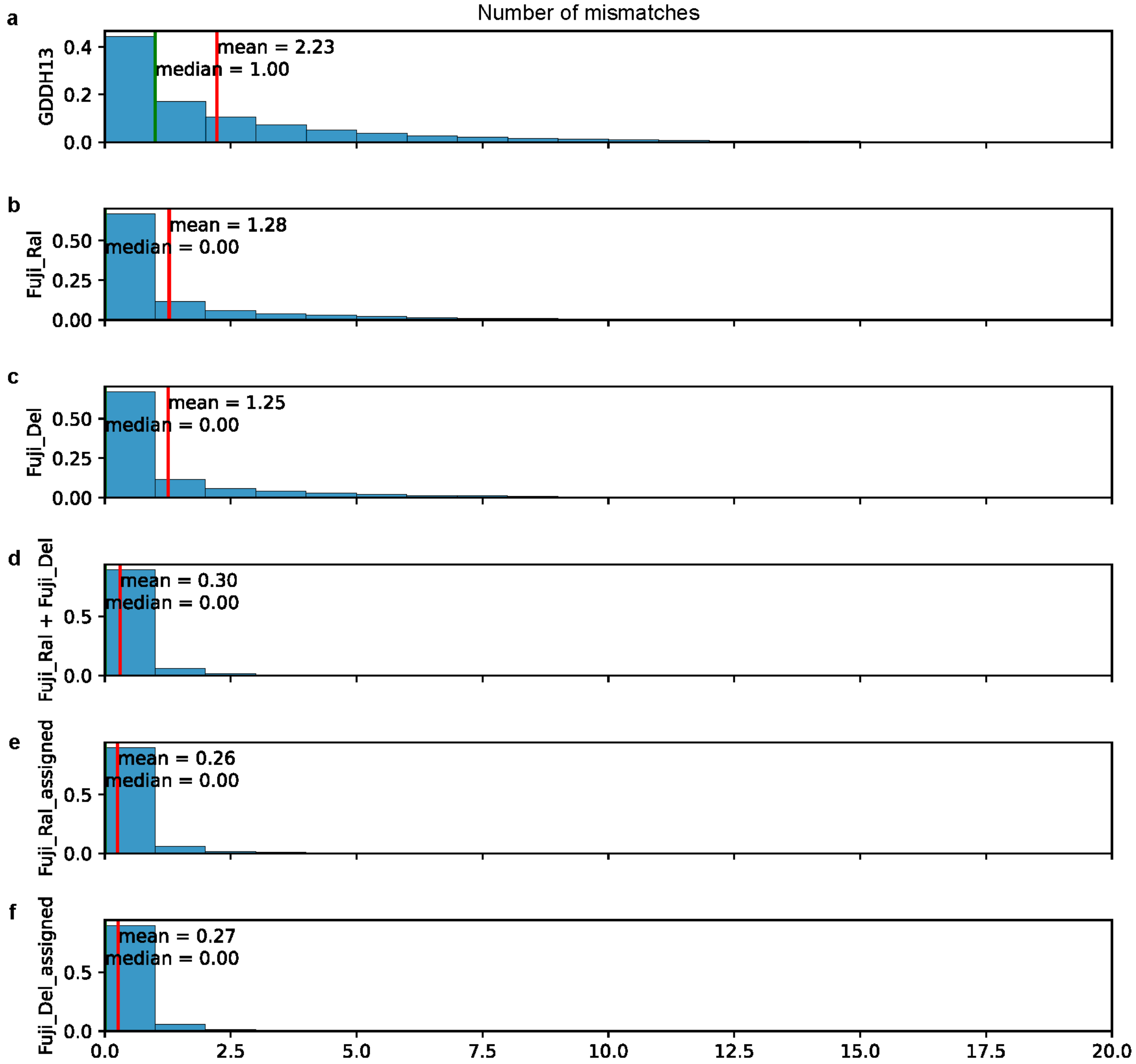


**Figure S13.** Number of mismatches using different references for read mapping. **a** Using GDDH13 as the reference genome. **b** Using Fuji_Ral as the reference genome. **c** Using Fuji_Del as the reference genome. **d** Using Fuji_Ral + Fuji_Del as the reference genomes without secondary assignment of reads. **e,f** Assigning reads to the appropriate haplotypes using the pipeline developed in this study.
